# Microtubule Lattice Spacing Governs Cohesive Envelope Formation of Tau Family Proteins

**DOI:** 10.1101/2021.11.08.467404

**Authors:** Valerie Siahaan, Ruensern Tan, Tereza Humhalova, Lenka Libusova, Samuel E. Lacey, Tracy Tan, Mariah Dacy, Kassandra M. Ori-McKenney, Richard J. McKenney, Marcus Braun, Zdenek Lansky

## Abstract

Tau is an intrinsically-disordered microtubule-associated protein (MAP) implicated in neurodegenerative disease. On microtubules, tau molecules segregate into two kinetically distinct phases, consisting of either independently diffusing molecules or interacting molecules that form cohesive “envelopes” around microtubules. Envelopes differentially regulate lattice accessibility for other MAPs, but the mechanism of envelope formation remains unclear. Here, we find that tau envelopes form cooperatively, locally altering the spacing of tubulin dimers within the microtubule lattice. Envelope formation compacted the underlying lattice, whereas lattice extension induced tau-envelope disassembly. Investigating other members of the tau-MAP family, we find MAP2 similarly forms envelopes governed by lattice-spacing, whereas MAP4 cannot. Envelopes differentially biased motor protein movement, suggesting that tau family members could spatially divide the microtubule surface into functionally distinct segments. We conclude that the interdependent allostery between lattice-spacing and cooperative envelope formation provides the molecular basis for spatial regulation of microtubule-based processes by tau and MAP2.

## Introduction

Microtubules, stiff polymers composed of tubulin dimers, are a key component of the cytoskeleton, providing mechanical rigidity for the cell and tracks for active intracellular transport driven by motor proteins. Microtubules undergo alternating phases of polymerization and depolymerization, which are dependent on the GTP hydrolysis activity of tubulin. During polymerization, GTP-tubulin associates with the growing tip of the microtubule. After tubulin incorporation into the microtubule lattice, tubulin-bound GTP is hydrolysed to GDP. The hydrolysis of GTP is accompanied by a ∼2.5Å longitudinal compaction of the interdimer interface within the mammalian tubulin lattice, reducing the lattice spacing of the microtubule (Alushin et al., 2014). However, it is important to note that lattice compaction upon GTP hydrolysis was not observed in yeast (Howes et al., 2017; von Loeffelholz et al., 2017) or worm (Chaaban et al., 2018) microtubule reconstructions, raising questions about the mechanism and conservation of this structural change within the lattice. Compacted GDP-lattices can be artificially extended by microtubule stabilizing agents, such as the GTP analogue Guanosine-5’-[(α,β)-methyleno]triphosphate (GMPCPP) or the anticancer drug taxol (Paclitaxel) (Alushin et al., 2014). While the slowly-hydrolyzable GMPCPP keeps the microtubule lattice spacing in a GTP-like extended state due to its extremely low rate of turnover (Hyman et al., 1992), taxol reversibly increases the microtubule lattice spacing, as the turnover of taxol binding to microtubules is high (Díaz et al., 2003). In addition to changes in the nucleotide state or the effect of drugs, tubulin length can also be altered by the interaction of microtubule-associated proteins (MAPs) with the microtubule lattice. For example, molecular motors of the kinesin-1 family or the tip-tracking protein EB3 alter microtubule lattice architecture (Morikawa et al., 2015; Peet et al., 2018; Zhang et al., 2018), demonstrating the dynamic interdependence between the structure of the microtubule lattice and lattice-bound proteins. MAPs have different affinities for the compacted or extended tubulin lattice and can employ this difference as a readout for their localization on the microtubule. Some MAPs have a higher affinity for the GTP-tubulin, which enables their specific localization to the microtubule tips, as is the case for the tip tracking proteins of the EB family (Maurer et al., 2011; Zanic et al., 2009) and some kinesin motors (S. Jiang et al., 2019; Zhernov et al., 2020). By contrast, other MAPs, such as the kinesin-3 Kif1a or the neuronal intrinsically disordered protein tau, are reported to have a higher affinity for GDP tubulin, which enables their specific localization to the GDP regions of the microtubules (Castle et al., 2020; Duan et al., 2017; Guedes-Dias et al., 2019; Tan et al., 2019).

Tau, along with MAP2 and MAP4, constitute a family of evolutionarily and structurally related, intrinsically disordered MAPs, which are especially important in neuronal development and function (Chapin & Bulinski, 1991; Dehmelt & Halpain, 2005; Sündermann et al., 2016). Tau is associated with a large number of neurodegenerative disorders collectively termed “tauopathies”, while, to date, any role of MAP2 and MAP4 in human disease is not described (Götz et al., 2019; Guo et al., 2017; Iqbal et al., 2005; Li et al., 2020; Strang et al., 2019). Tau can regulate the interactions of other MAPs with the microtubule surface by forming cohesive envelopes (previously referred-to as “condensates” or “islands”) which we define as spatial accumulations of tau molecules interacting with each other and the microtubule lattice. Tau envelopes effectively act as selectively permeable barriers for other MAPs (Siahaan et al., 2019; Tan et al., 2019). Consistent with this idea, tau mislocalization leaves axonal microtubules unprotected against microtubule-severing enzymes such as katanin (Qiang et al., 2006), leading to microtubule destabilization and the eventual degeneration of the axon. Tau may also be a key regulator of microtubule-based cargo trafficking in neurons. Tau directly inhibits the anterograde motility of kinesin-1 and kinesin-3 while allowing dynein-based retrograde motility (Chaudhary et al., 2018; Dixit et al., 2008; Ebneth et al., 1998; Monroy et al., 2020; Seitz et al., 2002; Siahaan et al., 2019; Tan et al., 2019; Trinczek et al., 1999; Vershinin et al., 2007). MAP2 is mostly localized to dendrites and the initial segment of the axon and regulates neuronal cargo trafficking by inhibiting kinesin motors (Gumy et al., 2017; Monroy et al., 2020; Seitz et al., 2002). In contrast to tau and MAP2, MAP4 is more ubiquitously expressed in various human tissues (Dehmelt & Halpain, 2005). Within neurons, MAP4 is localized in the axons, where it concentrates at axonal branching points (Tokuraku et al., 2010). MAP4 inhibits the association of the microtubule-severing enzyme katanin with microtubules (McNally et al., 2002) and has been reported to regulate microtubule-based cargo trafficking by inhibiting dynein motility and enhancing kinesin-based transport. However, a MAP4 fragment reduced the motility of kinesin-1 and kinesin-3 (Karasmanis et al., 2018; Samora et al., 2011; Semenova et al., 2014) *in vitro*, and overexpression of MAP4 largely inhibited vesicle motility *in vivo* (Bulinski et al., 1997).

Tau can assemble into various types of higher-order oligomers. In solution, tau can undergo liquid-liquid phase separation, and phase-separated tau condensates can nucleate new microtubules by concentrating tubulin from solution (Hernández-Vega et al., 2017). Tau condensates can also initiate another form of tau oligomer, tau aggregation (Hernández-Vega et al., 2017; Wegmann et al., 2018). In neurons, tau aggregation is strongly correlated with hyper-phosphorylation, and tau aggregates build up into very large neurofibrillary tangles (Iqbal et al., 2016), commonly found in the brains of Alzheimer’s disease patients. When bound to microtubules tau molecules can form oligomers (Dixit et al., 2008; McVicker et al., 2014) and cohesive superstructures which envelope the microtubule surface and regulate access to the underlying microtubule lattice (Siahaan et al., 2019; Tan et al., 2019). While cooperative binding to the microtubule surface has been reported for other MAPs (Bechstedt & Brouhard, 2012), the underlying mechanisms remain elusive. In particular, it is not known whether cooperative binding to microtubules arises solely due to direct interactions between the MAP molecules or if conformational changes of the underlying microtubule lattice are involved .

Here, we found that the spacing of tubulin dimers within the microtubule lattice governs the cooperative formation of cohesive tau envelopes on microtubules. At the same time, cooperative formation of tau envelopes reversibly affects the microtubule lattice spacing. We show that this mechanism is conserved in MAP2, but surprisingly, we find that the binding of MAP4 appears independent of the interdimer spacing within the microtubule lattice. We demonstrate distinct functional consequences of these differential binding modes within this family of evolutionarily related MAPs. Our results show that the regulation of MAP cooperativity through the spacing of tubulin dimers within the microtubule lattice spacing is an divergent evolutionary feature within this clinically important family of intrinsically disordered MAPs.

## Results

### Tau cooperativity through local microtubule lattice compaction

To study the formation of tau envelopes on the microtubule lattice, we immobilized taxol-stabilized microtubules on a coverslip, added purified full-length 2N4R tau (longest isoform), labeled with a fluorescent protein, and performed time-lapse imaging using total internal reflection fluorescence (TIRF) microscopy (Fig. 1a, Methods). Initially we varied the tau concentration from 2.6 nM to 100 nM and monitored the tau density on the microtubule surface. Quantifications of tau densities across this concentration range revealed that the formation of tau envelopes is a cooperative process (Fig. 1b, Methods) with a Hill coefficient of 1.9 ± 0.7. Intriguingly, during envelope growth we often observed that microtubules, which initially had a bent structure, straightened out when the envelope formed in the curved region (Fig. 1c, Movie S1), suggesting that cooperative binding of tau might locally influence the structure of the underlying microtubule lattice. Previously, we observed that tau envelopes readily form on GDP microtubule lattices but do not form on microtubules stabilized with GMPCPP (Tan et al., 2019). To understand how the structure of the microtubule lattice affects the formation of tau envelopes, we measured the growth rate of tau envelopes on different types of microtubule lattices by adding tau to (a) GDP-lattice microtubules (natively compacted), (b) taxol-lattice microtubules (extended reversibly), and (c) GMPCPP-lattice microtubules (extended irreversibly). We observed that tau envelope growth velocities on the natively compacted GDP-lattice microtubules (1.6 ± 1.3 μm/s; mean ± s.d) was faster by more than an order of magnitude than the growth velocity on the reversibly extended taxol-lattice microtubules (23.0 ± 12.8 nm/s; mean ± s.d), while, in agreement with previous observations (Tan et al., 2019), no envelope formation was observed on irreversibly extended GMPCPP-lattice microtubules (Fig. 1d, Supplementary Fig. 1a,b,c,d). These results demonstrate that tau envelopes have a strong preference to form on inherently compacted GDP-lattice microtubules.

**Figure 1:**
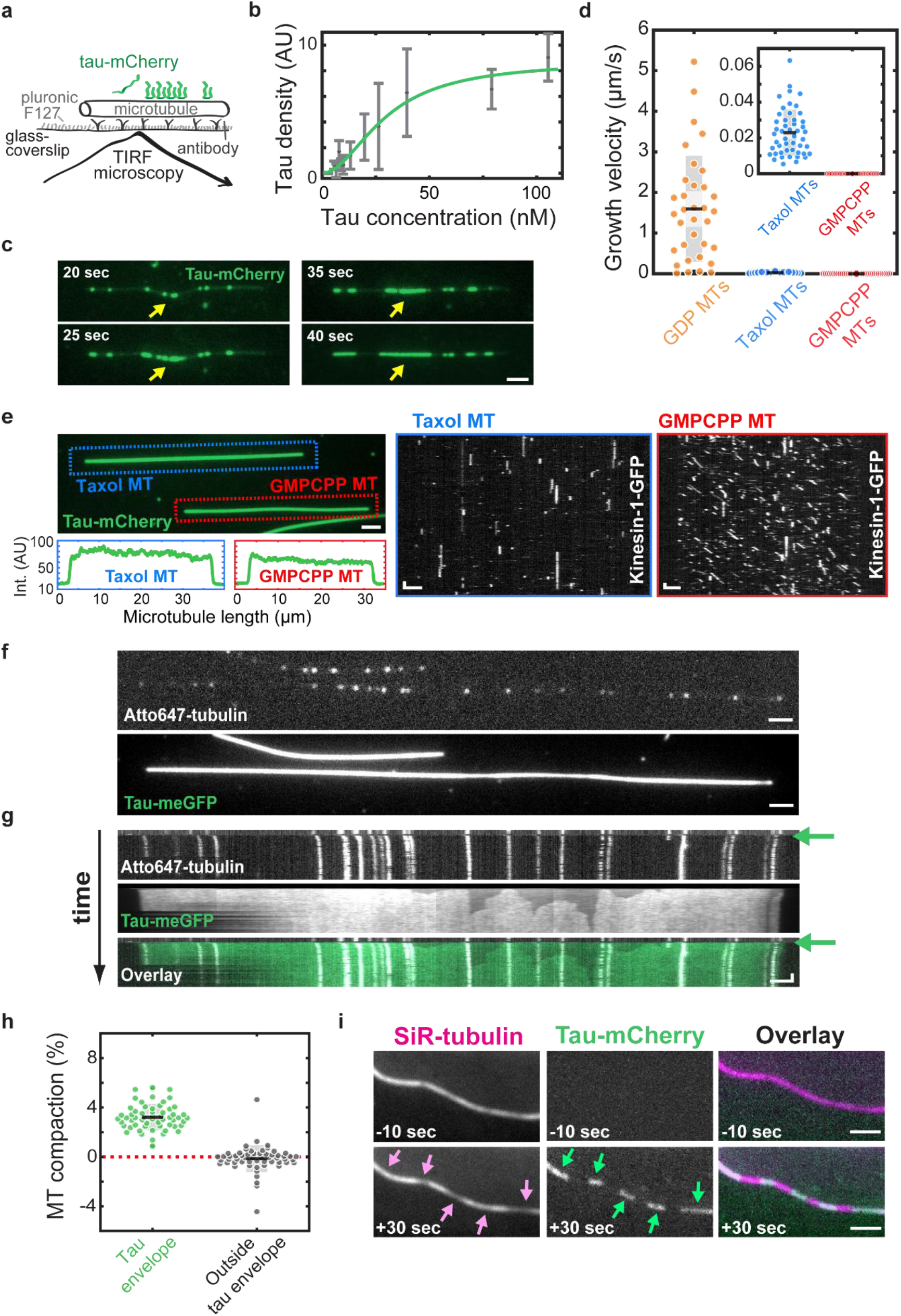
Tau cooperativity through local microtubule lattice compaction. **a.** Schematics depicting the assay geometry. **b.** Quantification of cooperative binding of tau to taxol-lattice microtubules. Data are mean ± s.d. (n=652 microtubules, 60 experiments, 95% confidence bounds, r-square = 0.9633, grey). Fit was done using Hill-Langmuir equation (green). **c.** Fluorescence time lapse micrographs showing changes in the microtubule lattice upon formation of tau-mCherry envelope (green). 20 nM tau-mCherry was added at t=0 sec. The yellow arrows indicate the position where the microtubule lattice straightens while the tau envelope grows. Scale bar: 2 μm. **d.** Growth velocities of tau envelopes on different microtubule lattices after the addition of 20 nM tau-mCherry. Growth velocity was 1.6 ± 1.3 μm/s (mean ± s.d., n=34 microtubules, 5 experiments) on GDP-lattice microtubules (orange), 23.0 ± 12.8 nm/s (mean ± s.d., n=49 microtubules, 3 experiments) on taxol-lattice microtubules (blue) while no envelope formation (n=200 microtubules, 14 experiments) was found on GMPCPP-lattice microtubules (red). **e.** Fluorescence micrograph (top left) showing a taxol-lattice microtubule (MT) (marked by a blue dotted rectangle) and a GMPCPP-lattice microtubule (marked by red dotted rectangle) after the addition of 600 nM tau-mCherry (green) and 60 nM kinesin-1-GFP. Fluorescence intensity profile along the microtubule length (bottom left) shows that the tau-mCherry density is comparable on both microtubules and envelopes are not distinguishable. The fluorescence kymographs (right panels) show that kinesin-1-GFP does not processively walk on taxol-lattice microtubules (left). By contrast, processive movement of kinesin-1-GFP was detected on GMPCPP-lattice microtubules (right). Scale bars: vertical 1s, horizontal 2 μm. **f.** Fluorescence micrograph of a speckled Atto647-labeled microtubule (top) immobilized on the coverslips surface after the addition of 400 nM tau-meGFP (bottom). Scale bar: 2 μm. **g.** Multichannel kymograph corresponding to the microtubule in **f**, showing compaction of the microtubule lattice by individual speckles (white) moving closer to each other after the addition of 400 nM tau-meGFP (green). The timepoint at which tau-meGFP was added to the microtubule is marked by green arrows. Scale bars: 2 μm, 1 min. **h.** Compaction of the microtubule lattice by tau within the envelope regions (tau envelope, green) and outside the envelope regions (outside tau envelope, grey). Outside of the envelope regions no compaction was found (-0.1 ± 1.1% compaction, mean ± s.d., n=57 regions, 7 experiments). **i.** Fluorescence micrographs of 2 μM SiR-tubulin (magenta) on a microtubule lattice before and after the addition of 20 nM tau-mCherry (green) at t=0 sec. The green arrows indicate the positions of the tau envelopes. The pink arrows indicate the corresponding local decrease in the SiR-tubulin density on the microtubule. Scale bar: 2 μm.

To test if irreversible lattice extension might completely prevent the formation of envelopes, even at saturating tau concentrations, we added 600 nM tau to a mixture of GMPCPP-lattice microtubules and compared them with taxol-lattice microtubules (Fig. 1e, left panels). At these elevated tau concentrations, tau uniformly covers the entire microtubule and the boundaries of envelopes cannot be distinguished by the local increase in tau density on the microtubule. However, the presence of tau envelopes can be determined by the effect they have on kinesin-1, which is occluded from microtubule segments enveloped by tau (Siahaan et al., 2019). After the addition of 60 nM kinesin-1 (in presence of 600 nM tau) to the microtubules, we observed no processive movement of kinesin-1 on the taxol-lattice microtubules, showing that these microtubules were fully covered by a tau envelope (Fig. 1e - right panels, Movie S2, for comparisons of kinesin-1 run lengths and landing rates see Supplementary Fig. 1e,f). By contrast, on the GMPCPP-lattice microtubules, processive movement of kinesin-1 motors was detected along their entire lengths, revealing that tau molecules on GMPCPP-lattice microtubules did not form tau envelopes, even at saturating tau concentrations. These results demonstrate that permanent extension of the microtubule lattice prevents envelope formation. Combined, these results suggest that the spacing of the microtubule lattice modulates the ability of tau to cooperatively bind to microtubules and thus regulates the formation of cohesive tau envelopes.

Therefore, we reasoned that tau envelope formation might induce microtubule lattice compaction on taxol-lattices that are extended reversibly. To test this hypothesis, we employed sparsely labeled microtubules (Fig. 1f, Supplementary Fig. 1g, Methods) containing fluorescent speckles to mark fixed positions within the microtubule lattice. These speckles provide a means to directly visualize and quantify microtubule lattice compaction upon cooperative binding of tau. After the addition of 400 nM tau to speckled taxol-lattice microtubules, we observed the rapid formation of a continuous tau envelope covering the entire microtubule lattice. Time-lapse imaging revealed that the relative distance between the individual speckles decreased during envelope formation, indicating that the microtubule lattice was compacting concomitantly with the cooperative binding of tau molecules (Fig. 1g, Movie S3). To determine whether the compaction is global (affecting the whole microtubule) or local (induced specifically in segments of the microtubule covered by the tau envelope), we added 20 nM tau to taxol-lattice speckled microtubules and followed the compaction in- and outside the envelope regions (Fig 1h, Supplementary Fig. 1g,h Methods). We found that there is no evident compaction of the lattice in the regions on the microtubule that were not covered by tau envelopes (57 non-envelope regions, 7 experiments), while the lattice regions that were covered by a tau envelope (on the same microtubule) showed a compaction of 3.2 ± 1.0% (mean ± s.d., 59 envelope regions, 7 experiments). Strikingly, the extent of the compaction closely matches the length difference between GTP- and GDP-bound tubulin dimers estimated previously by cryo-EM to be 2.4% (Alushin et al., 2014), suggesting that tau envelope shifts tubulin from its extended GTP-like into its compacted GDP-conformation. As cooperative binding of tau to the microtubule lattice reverses the taxol-induced lattice extension, we asked if the envelope formation will locally induce dissociation of taxol from the microtubule lattice. We therefore visualized the interaction between microtubules and the fluorogenic taxane SiR-tubulin. After incubating surface attached SiR-tubulin-lattice microtubules with 20 nM tau for 30 seconds we compared the SiR-tubulin density in the tau envelope regions to the density in the adjacent regions not covered by tau envelope (Fig. 1i, Movie S4). We detected a local decrease of 21.7 ± 12.9% in SiR-tubulin density in the envelope regions (mean ± s.d., n=72 envelopes, 5 experiments) showing that the tau-driven compaction of the microtubule lattice induces local dissociation of SiR-tubulin from the microtubule. These results are consistent with prior data showing competition between tau and taxol for microtubules (Kar et al., 2003). Combined these experiments show that the local structural shift to the compacted GDP-tubulin microtubule lattice induces the cooperative binding of tau molecules and is required for the cohesion of tau envelopes, and thus for the envelope formation and functionality.

### Lattice extension induces disassembly of tau envelopes

If tau envelope formation induces a reversible compaction of artificially (taxol-) extended tubulin, we hypothesized that physically lengthening the microtubule lattice should induce the disassembly of tau envelopes. To test this hypothesis, we extended the lattice by external force either locally or globally. Locally, we extended the microtubule lattice by bending the microtubules in a hydrodynamic flow. To do that, we used a low density of attachment points to immobilize GDP-lattice microtubules to the coverslip resulting in microtubules attached to the coverslip at a single point (Methods). After the addition of 20 nM tau to the microtubules, tau envelopes formed over the entire length of the GDP-lattice microtubules. By briefly (∼10 s) inducing a hydrodynamic flow in the experimental channel, the microtubules bent at their single attachment points, extending the lattice locally on the outside of the bent microtubule (Fig. 2a). During the hydrodynamic flow, we removed tau from solution and either added 10 μM taxol to induce subsequent lattice extension or kept the measurement buffer taxol-free. After the brief period of flow, the microtubules relaxed back to their original (straight) shape. We monitored the tau density at the attachment point during the whole experiment, i.e. before, during and after the period of flow. During the flow we found that, both in presence and absence of taxol in solution, the tau densities in the highly curved regions dropped to roughly 50% of their original density before hydrodynamic flow (Fig. 2b), indicating that the envelopes fissured at the positions of bending. After the hydrodynamic flow, in the presence of 10 μM taxol in solution, the envelopes disassembled from the ends. Additionally, the envelope started disassembling from the location where the fissure had been introduced by mechanical bending (Fig. 2a, Movie S6) as evidenced by the tau density drastically dropping to 3.6 ± 2.8% of the original value (Fig. 2a,b; mean ± s.d., n = 11 microtubules, 5 experiments). By contrast, in the absence of taxol, within a 200 s timeframe after the flow, the fissure closed when the microtubule relaxed and the envelope reformed, with the tau density recovering to 77.3 ± 10.8% (Fig. 2a,b, Movie S5; mean ± s.d., n=11 microtubules, 4 experiments). These results demonstrate that transient local expansion of the lattice by mechanical means destabilizes the tau envelope. Combining multiple external factors unfavorable to the existence of the tau envelope, such as physical bending of the microtubule and the addition of taxol, can disassemble the tau envelope completely. Thus, tau envelope formation and positioning are regulated both biochemically and physically.

**Figure 2:**
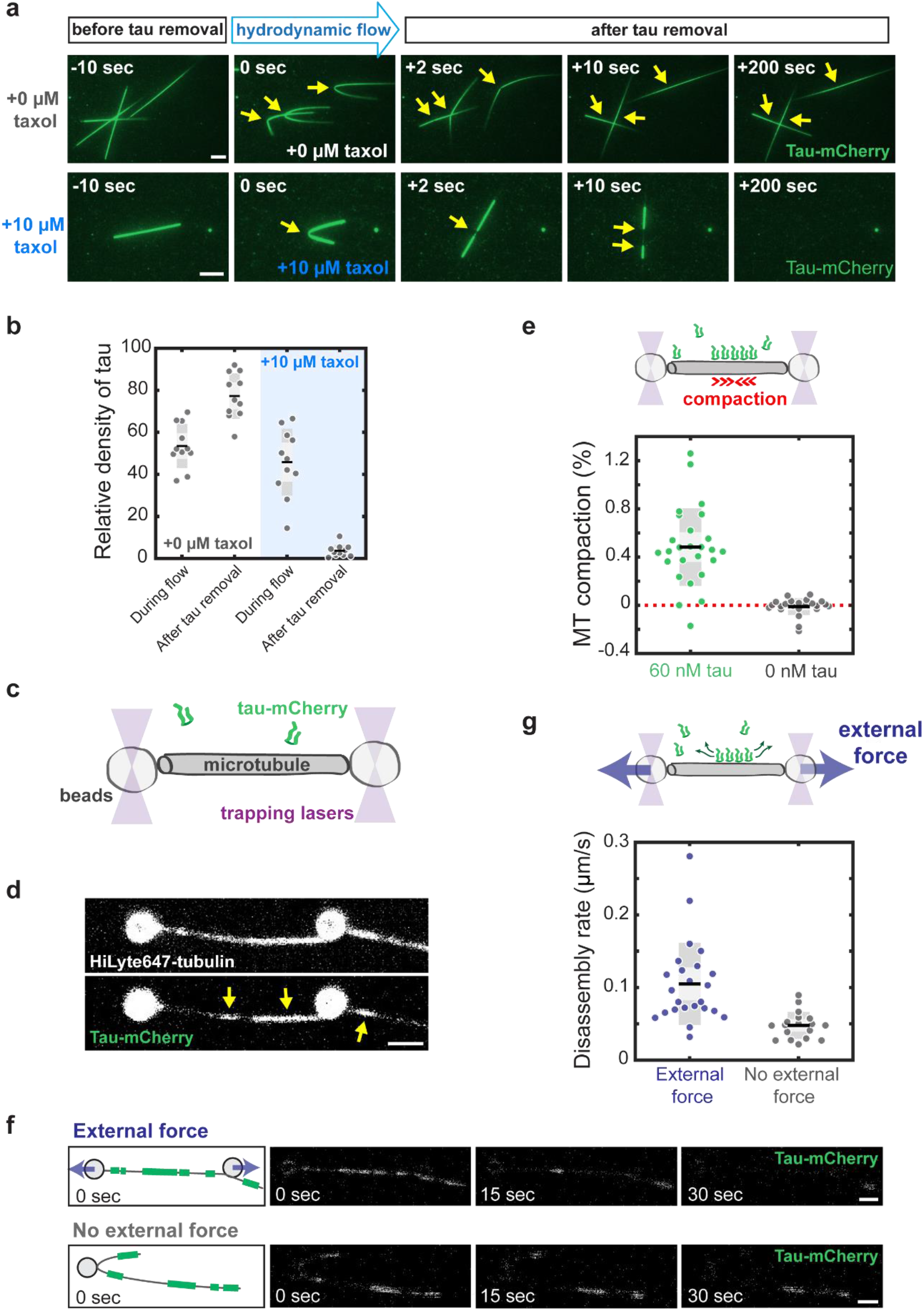
Lattice expansion induces disassembly of tau envelopes. **a.** Fluorescence micrographs of tau-mCherry envelopes on GDP-lattice microtubules before, during, and after removal of tau from solution at t=0 sec. During the hydrodynamic flow we either kept the measurement buffer taxol-free (top panels) or added 10 µM taxol (bottom panels). The yellow arrows indicate the location of the bend induced by the flow and the local decrease in the density of tau at the bent position. In absence of taxol, tau envelopes reformed and the density of tau in the bend recovered back to roughly 80% of its initial density. With taxol in solution, tau envelopes disassembled from their boundaries. Scale bars: 5μm. **b.** Relative density of tau-mCherry in the bend of GDP-lattice microtubules, during and after removal of tau-mCherry from solution. Intensities were measured during experiments either without taxol in solution (0 µM, left) or with 10 µM taxol in solution (10 µM, right) and normalized to the density before the brief period of hydrodynamic flow. Without taxol in solution the density dropped to 53.5 ± 10.4% during hydrodynamic flow (mean ± s.d.; n=11 microtubules, 4 experiments) and recovered back to 77.3 ± 10.8% of the original density (mean ± s.d.; n=11 microtubules, 4 experiments). With taxol in solution the density dropped to 45.9 ± 15.9% during hydrodynamic flow (mean ± s.d., n = 11 microtubules, 5 experiments) and decreased further to 3.6 ± 2.8% of the original density (mean ± s.d., n = 11 microtubules, 5 experiments). **c.** Schematics of the optical tweezers assay. A fluorescently labeled biotin-labeled taxol-lattice microtubule is suspended between two streptavidin-coated silica beads in a chamber containing tau-mCherry (green). **d.** Fluorescence micrographs of a biotin-HiLyte647-labeled taxol-lattice microtubule suspended between two beads after addition of 60 nM tau-mCherry. The yellow arrows indicate the locations of the tau envelopes. Scale bar: 2 μm. **e.** Compaction of a taxol-lattice microtubule measured after the addition of either 60 nM tau-mCherry (green) or 0 nM tau-mCherry (grey). With tau in solution the microtubule compacted 0.48 ± 0.32% (mean ± s.d., n = 26 microtubules). Without tau in solution the microtubule did not compact; 0.01 ± 0.07% compaction (mean ± s.d., n = 23 microtubules, 23 experiments, t-test, p= 3.85E-09). **f.** Fluorescence micrographs of a tau envelope disassembly experiment where the microtubule is stretched using external force (top panels) or relaxed in absence of external force (bottom panels). Sketches of the first micrographs (left panels) indicate the positions of the tau envelopes (green lines) at t=0 sec. Tau envelopes disassemble faster on microtubules that are stretched using external force. Scale bars: 2 μm. **g.** Disassembly rate of the tau envelopes on taxol-lattice microtubules either stretched by an external force or relaxed when no external force is applied. On stretched microtubules the envelope disassembly rate was 0.11 ± 0.06 μm/s (mean ± s.d., n = 24 microtubules, 24 experiments). On relaxed microtubules the envelope disassembly rate decreased to 0.05 ± 0.02 μm/s (mean ± s.d., n = 18 microtubules, 18 experiments, t-test, p= 0.0370).

To assess if global physical extension of the microtubule lattice can induce tau envelope disassembly, we specifically attached individual microtubules between two beads and stretched them using optical tweezers to uniformly extend the microtubule lattice. After suspending a taxol-lattice microtubule between two silica beads, we moved the beads slowly apart until we began detecting an increase in the force acting on the beads. We then fixed the beads in this position, with the microtubule in a straight but non-stretched state (Fig. 2c, Methods). We then added 60 nM tau to the chamber and observed tau envelope formation on the microtubule (Fig. 2d) and, simultaneously, detected an increase in force due to a decrease in the distance between the two beads, indicating a compaction of the microtubule lattice (Fig. 2e, Supplementary Fig. 2a). No compaction was detected in a control experiment when the buffer was exchanged but no tau was added (Fig. 2e, Supplementary Fig. 2b, n = 23 microtubules, 23 experiments). These results demonstrate that, on microtubule lattices previously extended reversibly through the administration of taxol, the formation of tau envelopes compact the extended lattices, and that this compaction generates significant forces as high as 40 pN.

Next, we stretched microtubules with pre-formed tau envelopes by applying an external force via slowly moving the beads apart until we reached a set force of 40 pN (Methods). We then removed tau from solution and monitored the disassembly rate of the tau envelopes, all while keeping the force at the constant value of 40 pN (Fig. 2f,g, Movie S7). In a control experiment, we monitored the disassembly rate of tau envelopes on microtubules suspended in a relaxed, non-stretched state (Movie S8). We found that the disassembly rate of tau envelopes increased by ∼ 2-fold when the microtubule lattice was stretched by an external force, indicating that global extension applied uniformly along microtubules, induces faster disassembly of tau envelopes. Combined, these experiments establish that tau envelopes impact microtubule mechanics and show that physically extending microtubule lattices coated by tau envelopes induces envelope disassembly, suggesting that tau functions in living cells may be mechanosensitive.

### Cooperative envelope formation is a divergent property within the tau-family

In vertebrates, the tau family also includes the primarily neuronal MAP2 and the more ubiquitously expressed MAP4. This family is characterized by high conservation within the carboxy-terminal microtubule binding repeats, but their sequences diverge significantly within their N-terminal and C-terminal projection domains (Fig. 3a) (Dehmelt & Halpain, 2005; Sündermann et al., 2016). All members of the family are expressed as multiple splice forms (Sündermann et al., 2016). Given our findings with tau so far, we wondered if cooperative lattice-gated envelope formation was conserved within the tau family. To investigate this question, we produced recombinant, fluorescently tagged MAP2c, the shortest isoform of MAP2, and the longest isoform of MAP4 (isoform 1) (Fig. 3a). We then examined the behavior of these MAPs on both taxol-lattice and GMPCPP-lattice microtubules (Fig. 3b). Strikingly, MAP2c behaved similarly to tau, forming obvious envelopes on taxol-lattice but not on GMPCPP-lattice microtubules. Conversely, we did not observe envelope formation on either lattice with MAP4 (Fig. 3b), but rather uniform binding along the microtubule length. We did not observe any evidence of envelope formation with MAP4 across a 10-fold range of concentrations on taxol-lattice microtubules (Supplementary Fig. 3a). Quantification of MAP fluorescence intensity on both types of lattices revealed that the level of MAP2c outside of the envelopes was identical to that on GMPCPP-lattice microtubules. This behavior is similar to tau (Tan et al., 2019). The intensity of MAP4 was identical on either type of lattice (Fig. 3b), further supporting the observation that MAP4 does not form envelopes. Thus, cooperative envelope formation appears to be a conserved feature of neuronally enriched tau and MAP2c, but not of the more ubiquitously expressed MAP4.

**Figure 3:**
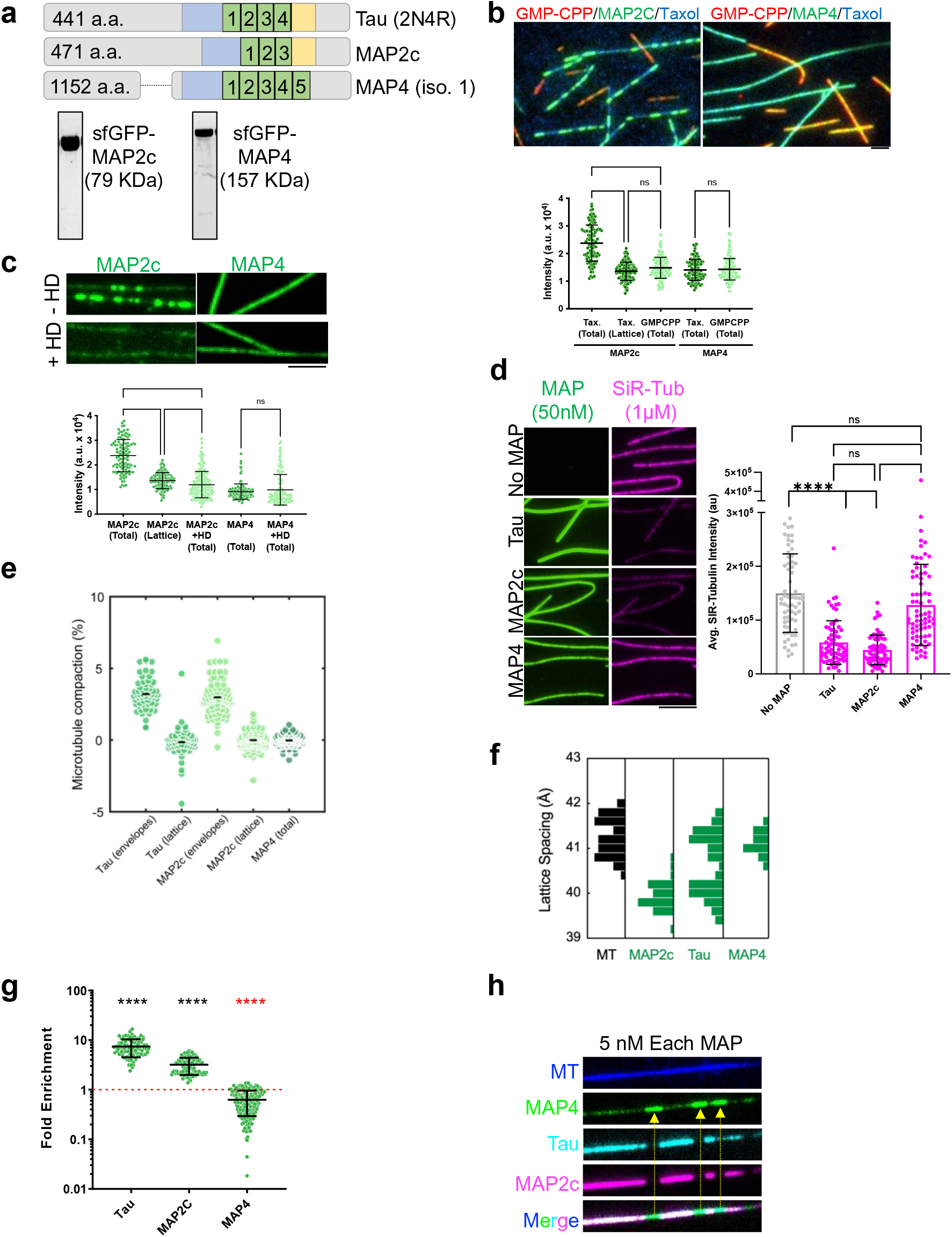
Cooperative envelope formation is a divergent property within the tau-family. **a.** Schematics of the MAP proteins analyzed, highlighting the conserved microtubule-binding regions (green), proline-rich region (blue), and pseudo-repeat (yellow). We utilized the longest splice form of tau (2N4R), the shortest splice form of MAP2 (isoform c), and isoform 1 of MAP4 containing 5 microtubule binding repeats. Below: Coomassie stained gels showing purity of the MAP2c and MAP4 proteins used. **b**. Multi-channel fluorescence micrographs showing the binding of 0.5 nM GFP-MAP2c or GFP-MAP4 (green) to either taxol-lattice (blue) or GMPCPP-lattice microtubules (red). Note the clear formation of envelopes by MAP2c on taxol-lattice, but not on GMPCPP-lattice microtubules. Below: quantification of the fluorescence intensity of MAPs on the indicated lattices. ‘Total’ refers to the intensity on the entire lattice including regions both outside and inside envelopes for MAP2c. N = 108, 134, 116, 106, 150 microtubule segments in 3 chambers each. Scale bar: 2 µm. **c.** Fluorescence images of 0.25 nM GFP-MAP proteins on taxol-lattice microtubules in the absence or presence of 10% 1,6-Hexanediol (HD). Below: quantification of the fluorescence intensity of MAPs in the indicated conditions. N = 108, 134, 199, 146, 125 microtubule segments respectively in 2 chambers each. Mean ± SD. **d.** Example fluorescence images showing GFP-MAP (green) and SiR-tubulin (magenta) signals along microtubules. Right: quantification of average SiR-tubulin fluorescence intensity in the absence or presence of the indicated MAPs. N = 65, 74, 72, 70 microtubules respectively in 2 chambers each. Mean ± SD. **e**. Compaction of the microtubule lattice measured on speckled microtubules after the addition of MAPs. Compaction of the microtubule lattice after the addition of 20 nM tau was 3.2 ± 1.0 % within the envelope regions (“envelope”, mean ± s.d., n= 59 envelopes, 7 experiments) and -0.1 ± 1.1 % outside the envelope regions (“lattice”, mean ± s.d., n= 63 lattices, 7 experiments). Compaction of the microtubule lattice after the addition of 1.5 nM MAP2c was 3.0 ± 1.2 % within the envelope regions (“envelope”, mean ± s.d., n= 78 envelopes, 5 experiments) and 0.0 ± 0.6 % outside the envelope regions (“lattice”, mean ± s.d., n= 95 lattices, 5 experiments). Compaction of the microtubule lattice after addition of 15 nM MAP4 was 0.0 ± 0.4 % over the entire length of the lattice (“total”, n = 105 microtubules, 11 experiments). **f.** Quantification of the tubulin monomer spacing from cryo-EM images of taxol-lattice microtubules in the absence or presence of the indicated MAPs. N = 46, 36, 44, 61, 30 microtubules analyzed respectively. **g.** Quantification of the enrichment of GFP-MAPs, based on fluorescence intensity, within mScarlet-2N4R tau envelopes. N = 97, 115, 94, 112, 206, and 183 tau envelopes, respectively from 2 chambers each. 0.5 nM MAP protein was used for each condition. Mean ± SD. Note that MAP4 values are below 1, meaning the protein is excluded from tau envelopes (red asterisks). **h**. Example image from multi-channel fluorescence experiment showing all three orthogonally-labeled MAPs mixed together on microtubules at equimolar concentration. Arrows denote the exclusion of MAP4 from the regions enriched with both tau and MAP2c. Scale bar: 5 µm.

To test whether the behavior of individual molecules constituting the envelopes is conserved within the tau family (Siahaan et al., 2019; Tan et al., 2019), we next examined the behavior of these MAPs on taxol-lattices at the single molecule level. At low picomolar concentrations, both MAP2c and MAP4 molecules showed predominantly diffusive interactions with microtubules (Supplementary Fig. 3a). To probe for any concentration dependent effects on single molecule behavior, we used higher concentrations of GFP-MAP2c (250 pM) or GFP-MAP4 (250 pM or 2.5 nM) and added low picomolar concentrations (10-30 pM) of TMR-labeled MAP2c or MAP4 that allow for the visualization of single molecules. The diffusive behavior of TMR-MAP2c outside of the GFP-MAP2c envelopes did not change with increasing concentrations of GFP-MAP2c. However, TMR-MAP2c molecules transitioned to largely static binding within the boundaries of the GFP-MAP2c envelopes (Supplementary Fig. 3a), where their dwell times increased ∼ 3-fold (Supplementary Fig. 3a,b): behavior very similar to that of tau (Siahaan et al., 2019; Tan et al., 2019). In contrast, TMR-MAP4 molecules remained largely diffusive regardless of the presence of higher concentrations of GFP-MAP4 (Supplementary Fig. 3a). Quantification revealed that the fraction of diffusive TMR-MAP4 molecules did not change over a 10-fold range of GFP-MAP4 concentrations (Supplementary Fig. 3a), consistent with an inability of MAP4 to form cohesive envelopes. To further explore the molecular dynamics of these MAPs on taxol-lattices, we performed FRAP experiments. Similar to previous observations with tau (Tan et al., 2019), MAP2c behaved differently inside the envelopes. MAP2c molecules turned over ∼ 4-fold faster when they resided outside versus inside envelope boundaries (Supplementary Fig. 3c). In contrast, MAP4 molecules showed homogenous behavior with a recovery time and mobile fraction more comparable to MAP2c molecules within envelopes, than out. This result is in accordance with the observed ∼2-fold longer dwell times for single MAP4 molecules versus MAP2c (Supplementary Fig. 3b). Tau envelopes are sensitive to the aliphatic alcohol 1,6-hexanediol (HD), a compound that disrupts phase separated systems in a variety of biological contexts (Tan et al., 2019), revealing that they share some material properties with these systems. Similar to tau (Tan et al., 2019), MAP2c envelopes dissolved when exposed to HD, but the diffusive fraction of MAP2c molecules remained unchanged (Fig. 3c, Supplementary Fig. 3b). In contrast, HD had no effect on the binding of MAP4 to microtubules, further showing the distinct biophysical properties of MAP4 (Fig. 3c, Supplementary Fig. 3b). These results reveal that tau and MAP2c envelopes share similar material properties that are distinct from MAP4. Combined, these results highlight distinctive behavior of MAP2c and MAP4 on the microtubule lattice and reveal closely related biophysical properties between tau and MAP2c.

Having established that tau envelope formation on the taxol-extended lattice leads to microtubule lattice compaction (Fig. 1), we wondered whether the same was true for MAP2c or MAP4. First, we mixed each MAP at high concentration (50 nM) with SiR-tubulin and measured the average SiR-tubulin intensity along microtubules. As previously noted (Fig. 1i), the presence of tau resulted in a decrease in SiR-tubulin intensity revealing tau-induced lattice compaction results in the removal of taxanes from the lattice (Fig. 3d). Similarly, the presence of MAP2c decreased SiR-tubulin binding by over 3-fold, while MAP4 had no effect (Fig. 3d). We next measured microtubule compaction by tracking speckled taxol-lattice microtubules upon addition of MAPs (Supplementary Fig. 1g,h). Similar to tau, we observed compaction of the speckle-labeled microtubule within MAP2c envelopes, but not outside (Fig. 3e). In contrast, we observed no microtubule compaction upon the addition of MAP4. We presume the decrease in SiR-tubulin fluorescence and compaction of microtubule speckles are due to conformational changes in the underlying tubulin lattice. To quantify this effect using a complementary method, we performed cryo-EM analysis of taxol-microtubules in the presence of MAPs to directly measure the tubulin spacing within the lattice. We found that both tau and MAP2c binding resulted in a decrease of the average spacing between tubulin monomers (Fig. 3f). Consistent with their ability to form envelopes that results in lattice reorganization, the addition of both tau and MAP2c caused a ∼1Å compaction between tubulin monomers, consistent with prior cryo-EM measurements between GDP and GTP-like lattices (Alushin et al., 2014). For tau, we observed a bimodal distribution of lattice spacing, suggesting that the lattice was not covered entirely in tau envelopes under our conditions. Addition of MAP2c caused a more complete compaction, to a level consistent with the compacted tau lattice (Fig. 3f). In contrast, addition of MAP4 did not result in measurable compaction within the tubulin lattice (Fig. 3f), consistent with the rest of our observations that MAP4 does not form cooperative envelopes on microtubules that alter the spacing of tubulin dimers within the lattice.

Given their distinct modes of binding, we wondered how the three members of the tau family co-exist on microtubules. We first mixed tau with either orthogonally-labeled tau, MAP2c or MAP4 and measured how enriched each molecule became within tau envelopes. As expected (Tan et al., 2019), tau was highly co-enriched within tau envelopes (Fig. 3g). Strikingly, MAP2c colocalized with the tau envelopes, where it was strongly enriched (Fig. 3g). By contrast, MAP4 was largely excluded from tau envelopes. To further confirm this result, we co-mixed all three MAPs at equimolar concentrations and observed strong co-segregation of tau and MAP2c into envelopes that largely excluded MAP4 into the regions surrounding the envelopes (Fig. 3h). These results highlight the strong similarities between tau and MAP2c and their biophysical differences with MAP4. They further reveal that tau and MAP2c form miscible MAP envelopes in vitro. Importantly, these experiments indicate that distinct properties of the microtubule lattice covered by cohesive envelopes provide a basis for the molecular selectivity for binding of orthogonal MAPs to the microtubule.

Taken together, these data reveal that like tau, MAP2c forms cohesive envelopes around microtubules, and these envelopes can compact an extended tubulin lattice. In stark contrast to tau and MAP2c, MAP4 does not form cohesive envelopes in any condition tested and is unable to affect the structure of the lattice to which it is bound. These results reveal previously unknown biophysical traits of this important family of MAPs and suggest functional diversification within the tau family could be related to the ability to recognize and alter the conformation of the microtubule lattice to which they are bound.

### Lattice spacing governs MAP cooperativity in vivo

As we have found that tau envelopes prefer microtubule lattices in a compacted GDP-bound state and lattice extension can induce disassembly of tau envelopes *in vitro*, we asked if we can induce tau envelope disassembly *in vivo* by extending the microtubule lattices in living cells with the addition of taxol. After the addition of 0.01 µM taxol to the culture medium, we imaged U-2 OS cells expressing tau for 10 minutes and monitored the tau density on the microtubules. In line with our *in vitro* observations, and previous *in vivo* evidence (Ettinger et al., 2016; Samsonov et al., 2004), we found that tau dissociated from the microtubules within the experimental timeframe (Supplementary Fig. 4a) as quantified by the decrease in the coefficient of variation of the tau signal measured before and 10 minutes after taxol treatment (Fig. 4a, Methods). In a control experiment, tau remained on the microtubules when DMSO was added to the medium in the absence of taxol (Supplementary Fig. 4a). Staining the cells for α-tubulin verified that microtubules were not disrupted by either treatment (Supplementary Fig. 4c,d). We repeated this experiment with U-2 OS cells expressing MAP4. In a strong contrast to tau and in line with previous *in vivo* evidence (Ettinger et al., 2016), we found that MAP4 remained on the microtubules after the taxol treatment (Supplementary Fig. 4b), as quantified by the unchanged coefficient of variation before and 10 minutes after the taxol treatment (Fig. 4a). Combined, these results show that tau unbinds from microtubules *in vivo* when the microtubule lattices are artificially extended by taxol, while MAP4 remains on the microtubules *in vivo* in the same conditions. These results are broadly consistent with our findings *in vitro* described above and show that tau is sensitive to the structure of the microtubule lattice in living cells.

**Figure 4:**
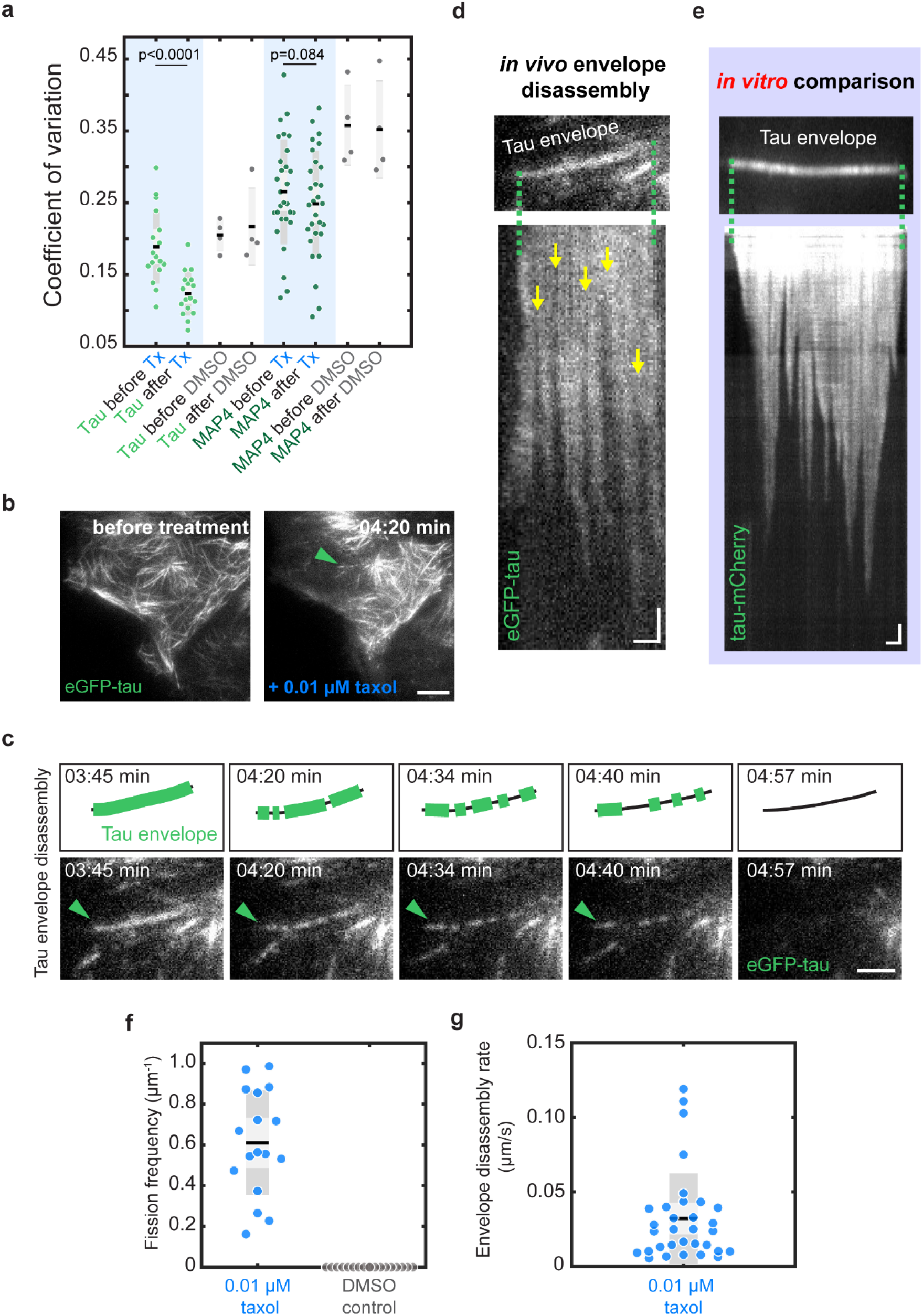
Lattice spacing governs MAP cooperativity in vivo. **a.** Coefficient of variation measured before and after taxol or DMSO treatment in eGFP-tau cells or eGFP-MAP4 cells. The coefficient of variation decreased significantly in cells overexpressing eGFP-tau (eGFP-tau before taxol treatment: 0.19 ± 0.05 (mean ± s.d.); eGFP-tau after taxol treatment: 0.12 ± 0.03 (mean ± s.d.), n = 17 cells, 5 experiments, t-test, p=4.548E-08), while the coefficient of variation in eGFP-MAP4 cells did not change significantly before and after taxol treatment (eGFP-MAP4 before taxol treatment: 0.27 ± 0.07 (mean ± s.d.); eGFP-MAP4 after taxol treatment: 0.25 ± 0.07 (mean ± s.d.), n = 29 cells, 5 experiments, t-test, p=0.084). During DMSO treatment (control) in eGFP-tau and eGFP-MAP4 cells, no significant difference in the coefficient of variation before and after DMSO addition was measured (eGFP-tau control before: 0.21 ± 0.02 (mean ± s.d.); eGFP-tau control after: 0.22 ± 0.05 (mean ± s.d.), n = 4 cells, 2 experiments, t-test, p=0.588; eGFP-MAP4 control before: 0.36 ± 0.06 (mean ± s.d.); eGFP-MAP4 control after: 0.35 ± 0.07 (mean ± s.d.), n=4 cells, 2 experiments, t-test, p=0.554,). **b.** Fluorescence micrographs showing that fissures appear in the eGFP-tau signal on the microtubules upon taxol treatment (green arrowhead). Scale bar: 10 μm. **c.** Time lapse fluorescence micrographs showing the progression of tau signal on the microtubule indicated in **b**. The fissures in the eGFP-tau signal grow in size and the tau envelope disassembles over 72 seconds. Scale bar: 2 μm. **d.** Kymograph of event shown in **b** and **c** showing the fissures (indicated by yellow arrows) appearing in the eGFP-tau signal and subsequent disassembly of the tau envelopes from their boundaries. Scale bars: horizontal 1 μm, vertical 10s. **e.** Kymograph of tau envelope disassembly *in vitro* showing striking resemblance with the *in vivo* observations in **e**. Scale bars: horizontal 2 μm, vertical 10s. **f.** Fission frequency of envelopes *in vivo* observed on microtubules covered with eGFP-tau after taxol treatment (0.01 μM taxol) or without the addition of taxol (control). Fission frequency observed after taxol treatment was 0.61 ± 0.26 μm^-1^ (mean ± s.d., n=17 microtubules in 8 cells). In the control experiment no fissions were observed (n=18 microtubules in 4 cells). **g.** Envelope disassembly rate after taxol treatment *in vivo* was 0.17 ± 0.31 μm/s (mean ± s.d., n=32 envelopes in 8 cells).

Importantly, when we analyzed the disassociation of tau from the microtubules during the taxol-driven extension of the compacted native microtubule lattice, we found that the tau signal disappeared from the microtubules in patterns that strikingly resembled the disassembly of tau envelopes *in vitro* (Siahaan et al., 2019; Tan et al., 2019). Without substantial changes in the initial tau density on the microtubule, fissures in the tau signal appeared, which increased in size over time. The distinct regions of the tau signal formed by these fissures then slowly disassembled from their boundaries until they fully disappeared (Fig. 4b-g, Movie S9,10). We interpret these findings as evidence for the existence of the cohesive tau envelope *in vivo*. The results suggest that native GDP-microtubule lattices *in vivo* can be fully enclosed by an envelope of cooperatively binding tau proteins, analogous to the envelopes observed *in vitro*.

### MAP cooperativity results in differential regulation of motor proteins

We next aimed to determine the functional implications for the different binding behaviors we observed for tau and MAP2c compared to MAP4. We therefore investigated how different concentrations of MAP2c and MAP4, spanning a 5-fold range, affect the motility of the retrograde microtubule motor complex, dynein-dynactin-Hook3 (DDH), and the anterograde kinesin-3 microtubule motor, KIF1A. DDH complexes contain two scaffolded dynein motors that work together to generate higher force and faster velocity along microtubules compared to complexes that contain a single dynein motor (Grotjahn et al., 2018; Urnavicius et al., 2018). In our assays, DDH complexes bound and moved processively along microtubules in the absence and presence of both MAP2c and MAP4 (Fig. 5a). We found that at the highest concentrations of MAPs tested, MAP4, but not MAP2c, decreased the number of processive dynein motors on the microtubule lattice (Fig. 5a,b). At low concentrations of MAP2c, where envelope boundaries are clearly visible, we did observe occasional pausing of DDH complexes at the envelope boundary (Fig. 5a, arrow), similar to our prior observations of tau’s effects on dynein-based transport (Tan et al., 2019). Neither MAP strongly affected the average velocity of these motors, although we measured a modest but statistically significant increase in DDH velocity at the highest concentration of MAP2c tested (Fig. 5c). We note that the average velocity measurements include the pauses induced by MAP2c envelopes and conclude the pausing behavior does not significantly affect the overall velocity of DDH motors. We analyzed the pixel-by-pixel correlation between the averaged intensities of the MAPs and DDH (methods). This analysis did not result in observable negative correlations between the average intensities of either MAP or the DDH motor complexes along the microtubule (Fig. 5d), suggesting neither MAP strongly affects the spatial distribution of the motors along microtubules. We note the modest effect of MAP4 on the DDH landing rate in our assays may be consistent with prior *in vivo* and *in vitro* results (Samora et al., 2011; Semenova et al., 2014), which suggested MAP4 could inhibit dynein-based movements. The landing rate of activated dynein-dynactin complexes is strongly determined by the binding of the p150^glued^ subunit of the dynactin complex (McKenney et al., 2016), and we therefore speculate that MAP4 may directly affect the interaction of p150^glued^ and the microtubule.

**Figure 5:**
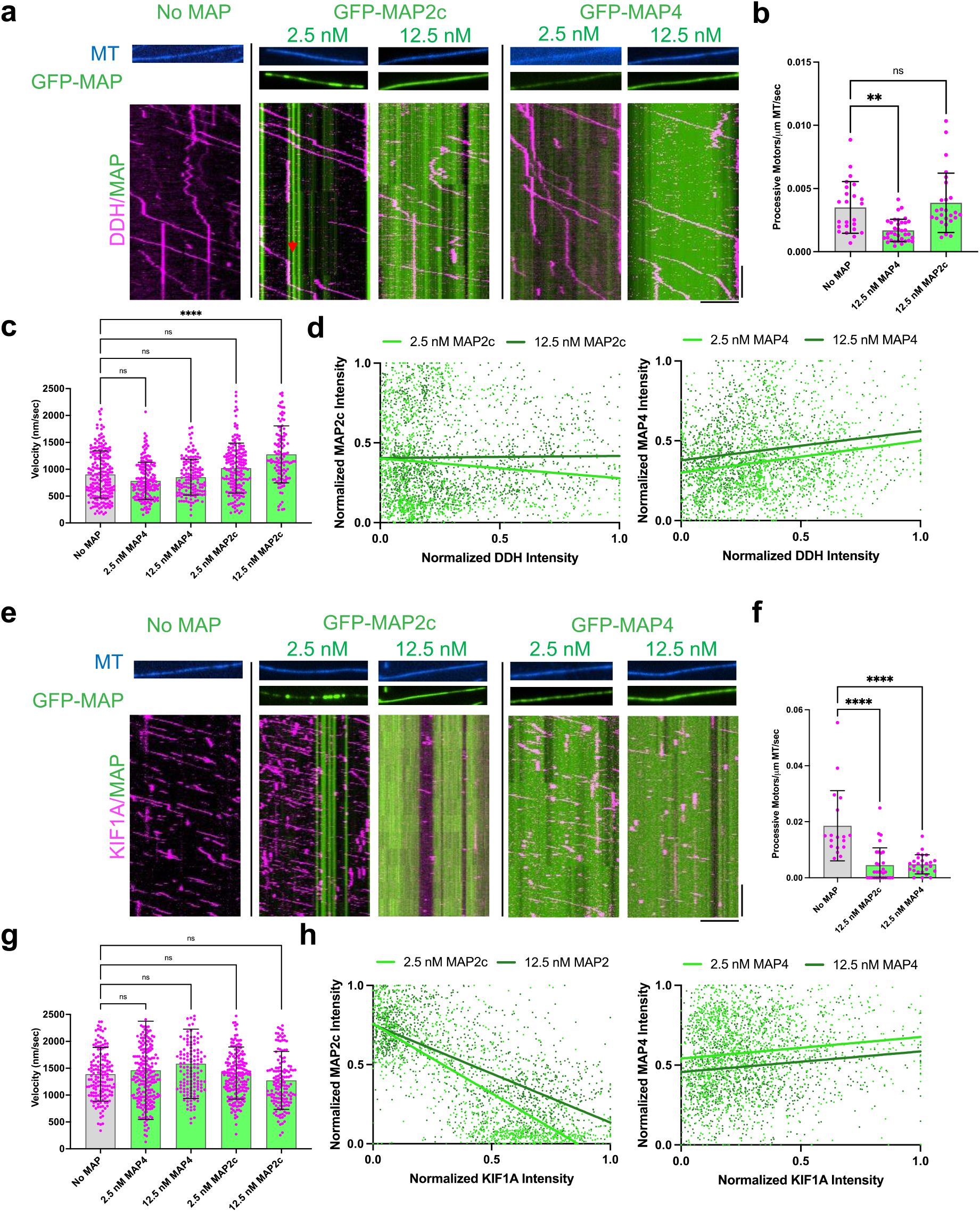
MAP cooperativity results in differential regulation of motor proteins. **a.** Representative example fluorescence micrographs showing microtubules (MT) (blue) and GFP-MAPs (green). Below each condition is a representative kymograph showing TMR-DDH (magenta) behavior. Red arrow denotes a pausing event. Scale bars: horizontal 5µm, vertical 10s. Note kymographs do not directly correlate with example images above. **b**. Quantification of the number of processive TMR-DDH complexes observed at the highest concentration of each GFP-MAP examined. Bars show averages with SD, magenta dots show individual measurement values: n = 24, 35, and 26 microtubules quantified from two independent trials each. * P=0.0014, ns- not significant by one-way ANOVA. **c.** Quantification of TMR-DDH velocities in the indicated conditions. Bars show averages with SD, magenta dots show individual measurement values: n = 201, 174, 143, 175, and 121 DDH complexes respectively, from N = 2 independent trials each. **** P<0.0001, ns- not significant by one-way ANOVA. **d.** Graphs displaying individual xy pairs per pixel for average GFP-MAP2c or GFP-MAP4 intensity versus average TMR-DDH intensity along the microtubule, fit with linear regression. Pearson’s correlation coefficients: -0.09669 and 0.01279 for 2.5 and 12.5 nM MAP2c respectively, and 0.1957 and 0.1671 for 2.5 and 12.5 nM MAP4 respectively. n= 1337 and 1557 xy pairs from n= 10 microtubules and N = two independent trials for 2.5 nM and 12.5 nM MAP2c. n = 1648 and 1193 xy pairs from n = 10 microtubules and N = two independent trials for 2.5 and 12.5 nM MAP4. P= 0.004, 0.6121, <0.001, and <0.001 for each 2.5, 12,5, 2.5, and 12.5 nM MAP2c and MAP4 respectively. **e.** Representative fluorescence micrographs showing microtubules (MT) (blue) and GFP-MAPs (green). Below each condition is a representative kymograph showing TMR-KIF1A (magenta) behavior. Scale bars: horizontal 5µm, vertical 10s. **f.** Quantification of the number of processive TMR-KIF1A motors observed at the highest concentration of each GFP-MAP examined. Bars show averages with SD, magenta dots show individual measurement values: n = 18, 26, and 30 microtubules quantified from two independent trials each. ****P<0.0001 by one-way ANOVA. **g.** Quantification of TMR-KIF1A velocities in the indicated conditions. Bars show averages with SD, magenta dots show individual measurement values: n = 158, 235, 138, 199, and 151 KIF1A motors respectively, from N = 2 independent trials each. ns-not significant by one-way ANOVA. **h.** Graphs displaying individual xy pairs per pixel for average GFP-MAP2c or GFP-MAP4 intensity versus average TMR-KIF1A intensity along the microtubule, fit with linear regression lines. Pearson’s correlation coefficients: -0.7620 and -0.6709 for 2.5 and 12.5 nM MAP2c respectively, and 0.1237 and 0.1201 for 2.5 and 12.5 nM MAP4, respectively. N = 1243 and 1354 xy pairs from n = 10 microtubules and N = two independent trials for 2.5 nM and 12.5 nM MAP2c. n = 1574 and 1121 xy pairs from n = 10 microtubules and N = two independent trials for 2.5 and 12.5 nM MAP4. P= <0.0001, <0.001, <0.0001, and <0.0001 for each 2.5, 12,5, 2.5, and 12.5 nM MAP2c and MAP4, respectively.

We also examined the processive kinesin-3 family member, KIF1A, which is known to transport cargo within neurons. Regulation of KIF1A transport within neurons is critical for human health as evidenced by a large cohort of human mutations in KIF1A that result in the neurodegenerative disorder KIF1A-associated neurological disease (KAND) (Boyle et al., 2021; Budaitis et al., 2021; Chiba et al., 2019; Lam et al., 2021). KIF1A has also been reported to be sensitive to the nucleotide state, and by extension possibly the dimer compaction state, of the microtubule lattice on which it travels (Guedes-Dias et al., 2019). We were thus interested to understand how KIF1A movement along microtubules within neurons may be influenced by tau-family MAPs that themselves affect the dimer spacing of the microtubule lattice. KIF1A motors bound and moved along microtubules in the absence and presence of low concentrations of MAP2c and MAP4 (Fig. 5e). However, we observed that higher concentrations of these MAPs strongly decreased the number of processive motors moving along the lattice without changing the velocity of those motors that were able to bind and move (Fig. 5e-g). These observations indicate that both MAPs strongly affect the landing rate of the KIF1A motors. In contrast to DDH, we observed a strong spatial effect of MAP2c on KIF1A (Fig. 5e,h). The average pixel intensities of KIF1A were highly negatively correlated with the average intensities of MAP2c at both concentrations of MAP tested, whereas we did not observe a negative correlation between KIF1A and MAP4 (Fig. 5h). These results indicate that cooperative envelope formation facilitates spatial regulation of the KIF1A motor along the microtubule and demonstrate that MAP2c and MAP4 exert different effects on retrograde and anterograde motors.

## Discussion

Cooperative binding of tau and MAP2c to microtubules both requires and, reciprocally, induces a local compaction of the underlying microtubule lattice. By contrast, the closely related MAP4 does not show this requirement and ability, demonstrating functional diversification within the tau family. What could be the mechanism underpinning cooperative envelope formation? We hypothesize two non-exclusive possibilities. We propose first that MAP-driven compaction of the lattice underlying the MAP envelope allosterically alters the conformation of the regions of lattice adjoining the boundaries of the envelope. This could be mediated by the conserved microtubule-binding repeats, which span several tubulin dimers with distinct binding properties (Kellogg et al., 2018). This “through the lattice” allostery might in turn locally increase the affinity for new MAP molecules binding from solution to the compacted lattice proximal to the enveloped boundaries. An allosteric mechanism operating through coupled changes in the microtubule lattice conformation has been previously proposed for the regulation of kinesin motors (Peet et al., 2018; Shima et al., 2018; Wijeratne et al., 2020) and microtubule dynamics (Kim & Rice, 2019). A second possibility is that direct MAP to MAP interactions might facilitate the preferential binding of new MAP molecules at the envelope boundaries. While it is currently unclear which regions of the MAPs may participate in such interactions, they could be mediated by the regions that flank the microtubule binding repeats of tau or MAP2, as these domains are required for envelope formation (Siahaan et al., 2019; Tan et al., 2019). Such interactions between MAP molecules would need to be sensitive to the conformational state of the underlying microtubule lattice, as we do not observe clear interactions between tau or MAP2 molecules on extended, GTP-like lattices. Both the N-terminal proline-rich domain and the C-terminal pseudorepeat domain are indispensable for tau envelope formation (Gustke et al., 1994; Siahaan et al., 2019; Tan et al., 2019). The pseudorepeat domain is conserved in MAP2 (61% identity) but not in MAP4 (14% identity), making this region a conspicuous target for future efforts to parse out the molecular mechanism of cooperative envelope formation.

What is the physical nature of cooperative MAP envelopes? Tau family MAPs are characterized as natively disordered proteins and tau itself undergoes liquid-liquid phase separation in solution and in cells (Hernández-Vega et al., 2017; Wegmann et al., 2018). Our results reveal that tau and MAP2c envelopes are sensitive to 1,6-hexanediol and thus share material properties with other liquid-liquid phase separated systems (Kroschwald et al., 2017; Shulga & Goldfarb, 2003). In addition, other MAPs such as TPX2 and BugZ, have been reported to undergo phase separation on the microtubule lattice (H. Jiang et al., 2015; King & Petry, 2020). We suggest that it is therefore conceivable that phase separation of tau and MAP2 on the microtubule surface might contribute to cooperative envelope formation.

We observed that MAP envelopes form on lattices reversibly extended by the addition of taxol, while they do not form on the permanently extended lattices in the presence of GMPCPP. We ascribe this difference is to the fast turnover of taxol and provide direct evidence for this hypothesis by demonstrating the displacement of taxanes from microtubule lattices by growing tau envelopes. Consistently, the very low hydrolysis rate of GMPCPP (Hyman et al., 1992), locking the microtubules in a permanently extended conformation, prevents tau and MAP2 from forming envelopes on this type of lattice, explaining a recent report of the preferential distribution of tau away from the growing plus-ends of dynamic microtubules (Castle et al., 2020).

Our data suggests that tau and MAP2, but not MAP4, are sensitive to the mechanical state of the microtubule lattice, and thus are suited to function as mechanosensitive MAPs. This observation may have physiological relevance in muscle cells where isoforms of MAP4 are abundantly expressed, and there are several reports of tau expression in various types of muscle tissue (Gu et al., 1996; Li et al., 2020; Shults et al., 2020; Trujillo et al., 2013). Additionally, mechanosensitivity could be an important feature in the developing nervous system during cell polarization or cell migration.

Structural studies on mammalian tubulin have revealed that the GDP lattice primarily adopts the compacted state, whereas the region near the growing plus-end that contains newly incorporated GTP tubulin adopts an expanded state (Alushin et al., 2014). In agreement with earlier findings (Tan et al., 2019), tau binding is disfavored in this region on dynamic microtubules (Castle et al., 2020). Intriguingly, in non-mammalian species, microtubule lattice compaction does not appear to be coupled to GTP hydrolysis, as structural studies have revealed that both yeast and worm microtubules have an extended GDP lattice (Chaaban et al., 2018; Howes et al., 2017; von Loeffelholz et al., 2017). These data raise questions about the compaction state of the microtubule lattice in living cells and suggest that the tubulin compaction state may not be as directly coupled to nucleotide hydrolysis as previously thought. Additionally, recent evidence suggests that after polymerization, the tubulin lattice is not conformationally homogenous, but rather releases and incorporates new GTP-tubulin subunits in response to mechanical damage and repair (Triclin et al., 2021; Vemu et al., 2018). Thus, the tubulin lattice in living cells may not exist in a homogenously compacted, GDP state, as previously thought. We suggest it is plausible that MAPs that recognize and alter the compaction state of the lattice, such as tau and MAP2, may play active roles in biasing the conformational dynamics of the tubulin lattice.

Given the structural data described above, it is presumed that the predominant volume of the microtubule lattice in cells is in the GDP compacted state, with GTP-tubulin present only at the short regions at tips of growing microtubules (Seetapun et al., 2012) and at sites of lattice damage. This distribution of GDP and GTP lattices suggests that most of tau in cells is bound to the microtubule in a cooperative manner. This notion agrees with our experimental observations *in vivo*, where most of the visible tau unbound from microtubules after the addition of taxol, and this unbinding occurred in a manner very similar to envelope dissolution *in vitro* (Siahaan et al., 2019). Initially, fissures formed in the uniform tau signal along the microtubules, forming envelopes of tau signal which decreased in size as more tau molecules dissociated from their boundaries. After the complete disassembly of these tau envelopes, we observed very little tau signal on straight microtubules. We hypothesize that as tau can only bind microtubules non-cooperatively after the taxol treatment, this non-cooperative interaction is too weak to be visualized in the high background signal from the dissociated tau in the cytoplasm. Another possibility is that tau’s association with taxol-lattice microtubules is weak, resulting in tau being outcompeted from the microtubules by other MAPs present in the cell (Monroy et al., 2020). This idea would also explain the relatively high sensitivity of tau’s localization on microtubules to low amounts of taxanes *in vivo*, as compared to *in vitro* (Fig. 3 and 4). However, on sparsely tau decorated microtubules, we occasionally observed an enriched tau signal in microtubule bends, even after taxol addition, in accordance with prior observations (Ettinger et al., 2016; Tan et al., 2019). This is likely due to tau binding to the compressed lattice on the inside of the bends. Furthermore, our results could provide a mechanistic explanation for observations showing that up-regulation of tau decreases sensitivity of cancer cells to taxol treatment (Rouzier et al., 2005; Smoter et al., 2011).

We have demonstrated that MAP envelopes differentially regulate the motility of two of the predominant microtubule motor systems in cells: dynein-based retrograde transport and kinesin-based anterograde transport. Our initial studies on tau (Siahaan et al., 2019; Tan et al., 2019) demonstrated disparate effects of tau envelopes on dynein versus kinesin-1 transport, with dynein largely able to pass through tau envelopes while kinesin-1 was completely excluded. We have extended this analysis here to MAP2 and MAP4 and found that formation of cooperative tau or MAP2 envelopes provides spatially distributed gating for the passage of anterograde KIF1A motors, but not retrograde dynein motors. Because MAP4 does not form envelopes, it is unable to provide spatially distinct regulation of motors and globally inhibits the landing of both dynein and kinesin onto microtubules. These data suggest that in the absence of extrinsic factors, MAP4 cannot act as a spatial regulator of microtubule-based trafficking in cells. However, our observation that MAP4 is also excluded from tau or MAP2 envelopes raises the possibility that MAP exclusion could act as one such extrinsic mechanism to spatially dictate MAP binding within cells. Such a mechanism has also been suggested for tau and MAP7, which have opposite effects on kinesin-1 motility (Monroy et al., 2020).

Why are different classes of kinesin motors unable to access the microtubule lattice under MAP envelopes? CryoEM structures suggests that kinesin and tau share partially overlapping binding sites on the lattice and thus a steric clash may prevent co-occupation (Kellogg et al., 2018; Tan et al., 2019). Additionally, kinesin binding is thought to cause the lattice to expand into the GTP like state (Peet et al., 2018). Therefore, it is possible that, rather than a direct steric clash between kinesin and MAPs, exclusive binding arises from competition for an expanded versus a compacted state of the microtubule lattice. The structure of MAP4 bound to microtubules demonstrates that it adopts a conformation completely distinct from that of tau (Kellogg et al., 2018; Shigematsu et al., 2018). This observation may at least partially explain why, unlike tau, MAP4 appears unable to form envelopes and does not exclude the binding of kinesin to microtubules like tau does. In support of this idea, previous lower resolution structures of tau and MAP2 bound to microtubules have demonstrated a strikingly similar structure for both of these MAPs (Al-Bassam et al., 2002) . In prior experiments, MAP4 did not block the binding of a kinesin-1 motor domain to microtubules (Shigematsu et al., 2018). However, our data show that it does negatively affect the ability of KIF1A to bind and move along microtubules, possibly revealing distinct effects on various classes of kinesin motors. This distinction could be due to the dominating influence of KIF1A’s k-loop on its landing rate (Soppina & Verhey, 2014), which may be particularly affected by the presence of MAP4. In sum, our data reveal that cooperative envelope assembly by tau and MAP2 spatially gates kinesin access to the underlying microtubule lattice, while allowing dynein to pass through these regions largely uninhibited. These MAPs are highly enriched within the neuronal system and it is tempting to speculate that this capability may be harnessed by highly elongated cells to spatially regulate microtubule-based trafficking. On the other hand, the more broadly expressed MAP4 may act to globally tune microtubule motor transport, but further work is necessary to fully understand if and how the tau family MAPs tune active transport within cells.

Tubulin lattice spacing regulates the affinity of microtubule associated proteins for the microtubule lattice. Here we have shown that protein binding, in particular envelope formation, locally changes microtubule lattice spacing and vice versa. Colocalization of tau and MAP2 into shared envelopes that exclude MAP4 suggests that different MAPs can form spatially distinct domains on the microtubule lattice, differentially regulating access to microtubule surface. Our work raises the hypothesis that the heterogeneity of the MAP envelope may thus provide a means for sectioning the microtubule surface into functionally distinct segments. We anticipate that the discovery of cooperative envelope formation provides new insights into the elusive physiological function, and possibly the disease mechanism, of the tau family of MAPs.

### Supplemental information

**Supplementary Figure 1 related to Figure 1.**
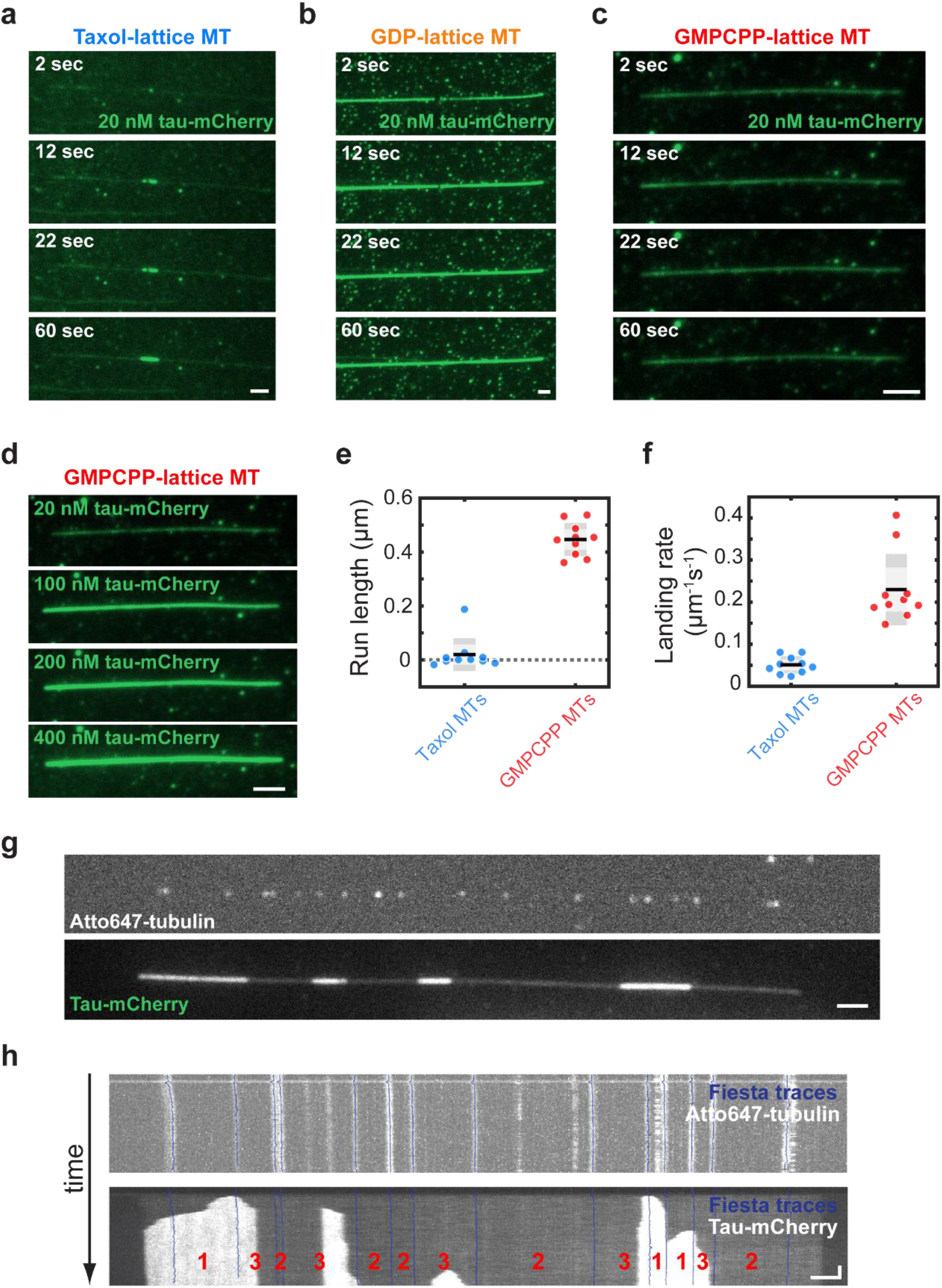
**a.** Tau envelopes growing on a taxol-lattice microtubule after the addition of 8 nM tau-mCherry. Scale bar: 2μm. **b.** Tau envelopes rapidly growing on a GDP-lattice microtubule after the addition of 8 nM tau-mCherry. Scale bar: 2μm. **c.** and **d.** No envelope formation on a GMPCPP-lattice microtubule after the addition of 8 nM tau-mCherry. In contrast to **a** and **b** the tau-mCherry density is uniform along the microtubule length over time and also at increasing concentrations. Scale bar: 2 μm. **e.** Run length of kinesin-1-GFP on taxol-lattice microtubules compared to GMPCPP-lattice microtubules in the presence of 600 nM tau-mCherry. Kinesin-1-GFP run length was 19.4 ± 60.3 nm/s on taxol-lattice microtubules (mean ± s.d., n=10 microtubules (281 molecules), 4 experiments). On GMPCPP-lattice microtubules the kinesin-1 run length was 446.7 ± 61.1 nm/s (mean ± s.d., n=10 microtubules (1058 molecules), 5 experiments). **f.** Landing rate of kinesin-1-GFP on taxol-lattice microtubules compared to GMPCPP-lattice microtubules in the presence of 600 nM tau-mCherry. Kinesin-1-GFP landing rate on taxol-lattice microtubules was 0.05 ± 0.02 μm^-1^s^-1^ (mean ± s.d., n=10 experiments (281 molecules), 4 experiments). Kinesin-1-GFP landing rate on GMPCPP-lattice microtubules was 0.23 ± 0.08 μm^-1^s^-1^ (mean ± s.d., n=10 microtubules (1058 molecules), 5 experiments) **g.** Fluorescent micrograph of an Atto647-labeled speckled microtubule (top) after addition of 20 nM tau-mCherry (bottom). Scale bars: 2μm, 1 min. **h.** Visual representation of compaction analysis using Fiesta tracking software. Fluorescent kymographs correspond to the microtubule from **d**. Individual speckles on the microtubule lattice can be traced with Fiesta tracking software (top) and using the fluorescent kymograph of the tau signal, areas between the traces are assigned the corresponding type of event (bottom). Event types are indicated by red numbers. 1: Tau envelope region, 2: Non-envelope region, 3: Not measured (both events occurred). Scale bar: 2μm.

**Supplementary Figure 2 related to Figure 2.**
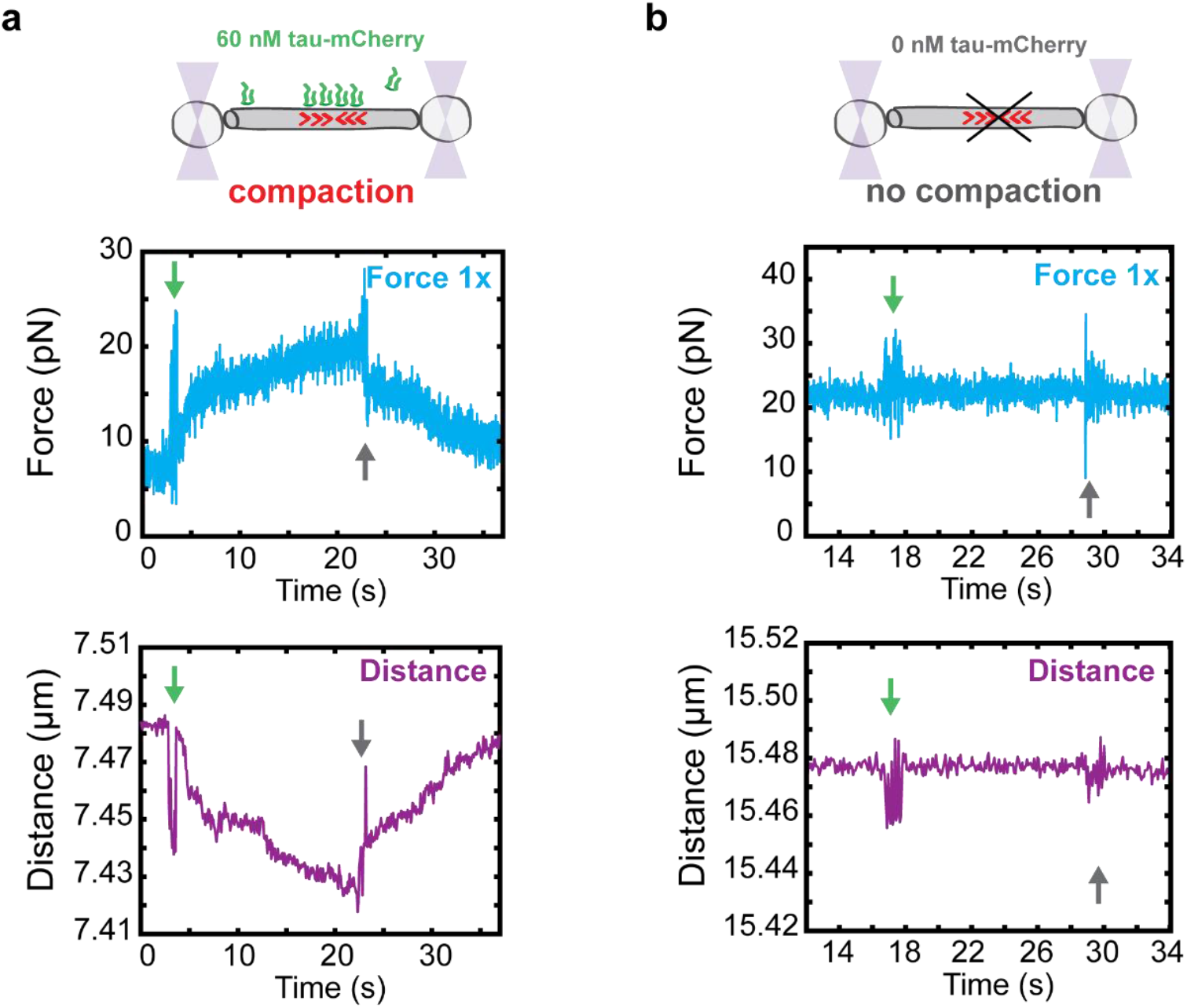
**a.** Representative force-time (blue) and distance-time (purple) graphs from a compaction experiment of a single taxol-lattice microtubule after addition of 60 nM tau-mCherry to the chamber. The green arrows indicate the timepoint at which 60 nM tau-mCherry was added. After this point the distance between the two beads decreased while the detected force increased, indicating compaction of the microtubule lattice. The grey arrows indicate the timepoint at which tau-mCherry was removed from the channel. After this point an increase in the distance and a decrease in the force is detected, indicating the relaxation of the microtubule lattice as tau unbound. **b.** Representative force-time (blue) and distance-time (purple) graphs from a control experiment where no tau-mCherry was added to the chamber. The green and grey arrows indicate the timepoints at which 0 nM tau-mCherry was added and removed, respectively. After these points no change in the force or in the distance was measured, indicating that no compaction or relaxation of the microtubule lattice occurred.

**Supplementary Figure 3 related to Figure 3:**
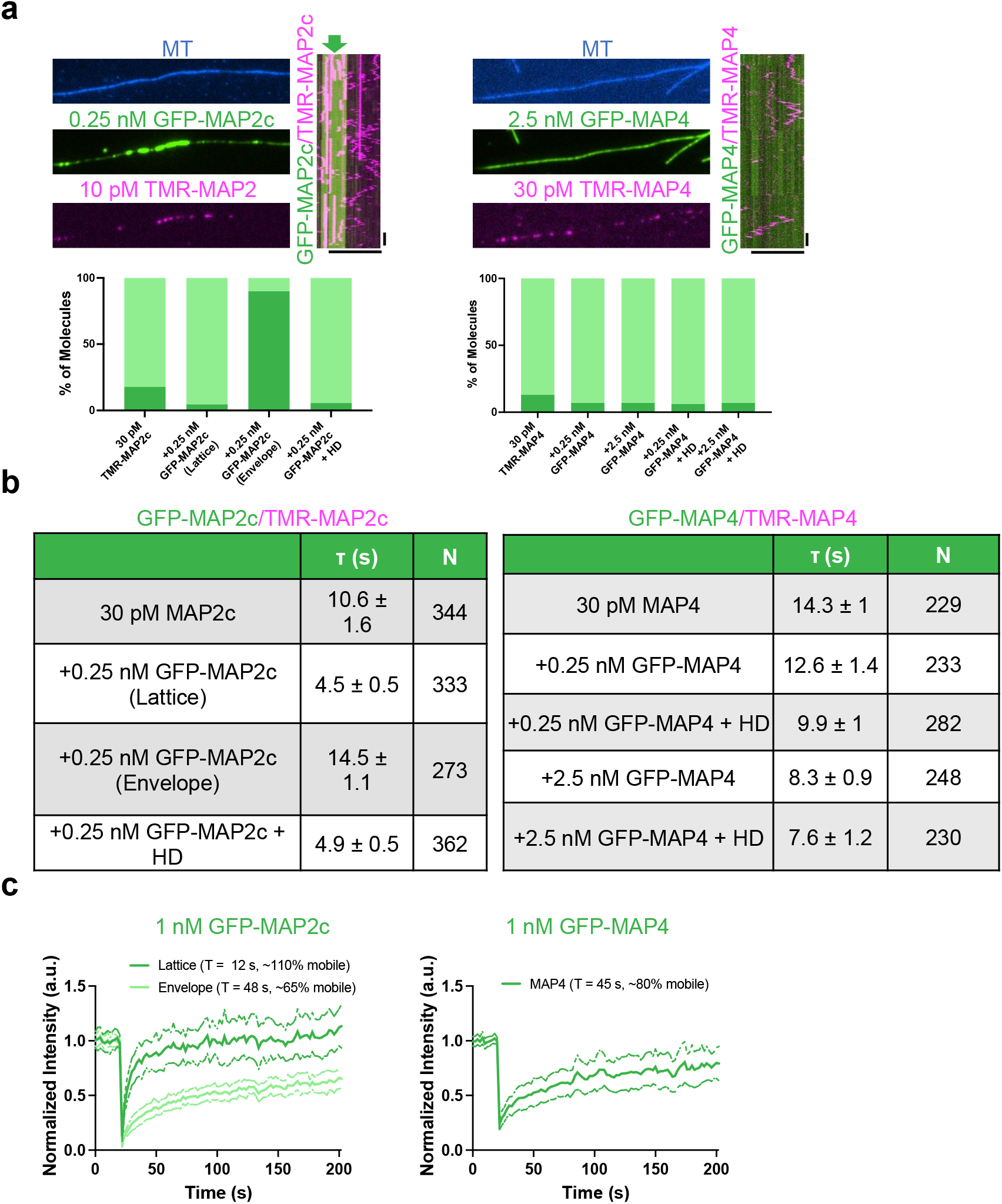
Characterization of MAP2c and MAP4 dynamics on taxol-lattice microtubules. **a.** Fluorescence micrographs from a multi-channel TIRF image showing the taxol-lattice microtubule (blue), higher concentration GFP-MAP (green) and single molecule concentration TMR-MAP (magenta). Right: kymographs showing the behavior of single TMR-MAP molecules in the presence of higher concentrations of GFP-labeled MAP. Note the majority of TMR-MAP2 molecules diffuse on the lattice outside of the GFP-MAP2 envelope (green arrow), but are statically bound inside the envelope. MAP4 molecules were largely diffusive regardless of the presence of higher concentrations of GFP-MAP4. Scale bars: horizontal 5µm, vertical 10s. Fractions of diffusive (light green bars) or statically bound (dark green bars) MAPs are shown below each panel. Note the large change in MAP2c behavior between inside and outside the envelopes. b. Table showing the quantifications of single molecule dwell times of TMR-MAP2c or TMR-MAP4 in the indicated conditions. Dwell times were fit to a single exponential decay function and the characteristic time constant (***t***) is reported as the dwell time with 95% CI errors of the fit shown. N = number of molecules quantified from at least 2 chambers each. c. Averaged FRAP recovery curves shown for each MAP. For MAP2c, FRAP was performed both outside and inside envelopes. The fitted recovery rate and fraction of mobile molecules is shown for each MAP. Bold lines show the average measured intensity and SD of the measurements. N = 91 measurements for each MAP condition including lattice vs envelopes for MAP2c from at least 2 chambers each.

**Supplementary Figure 4 related to Figure 4.**
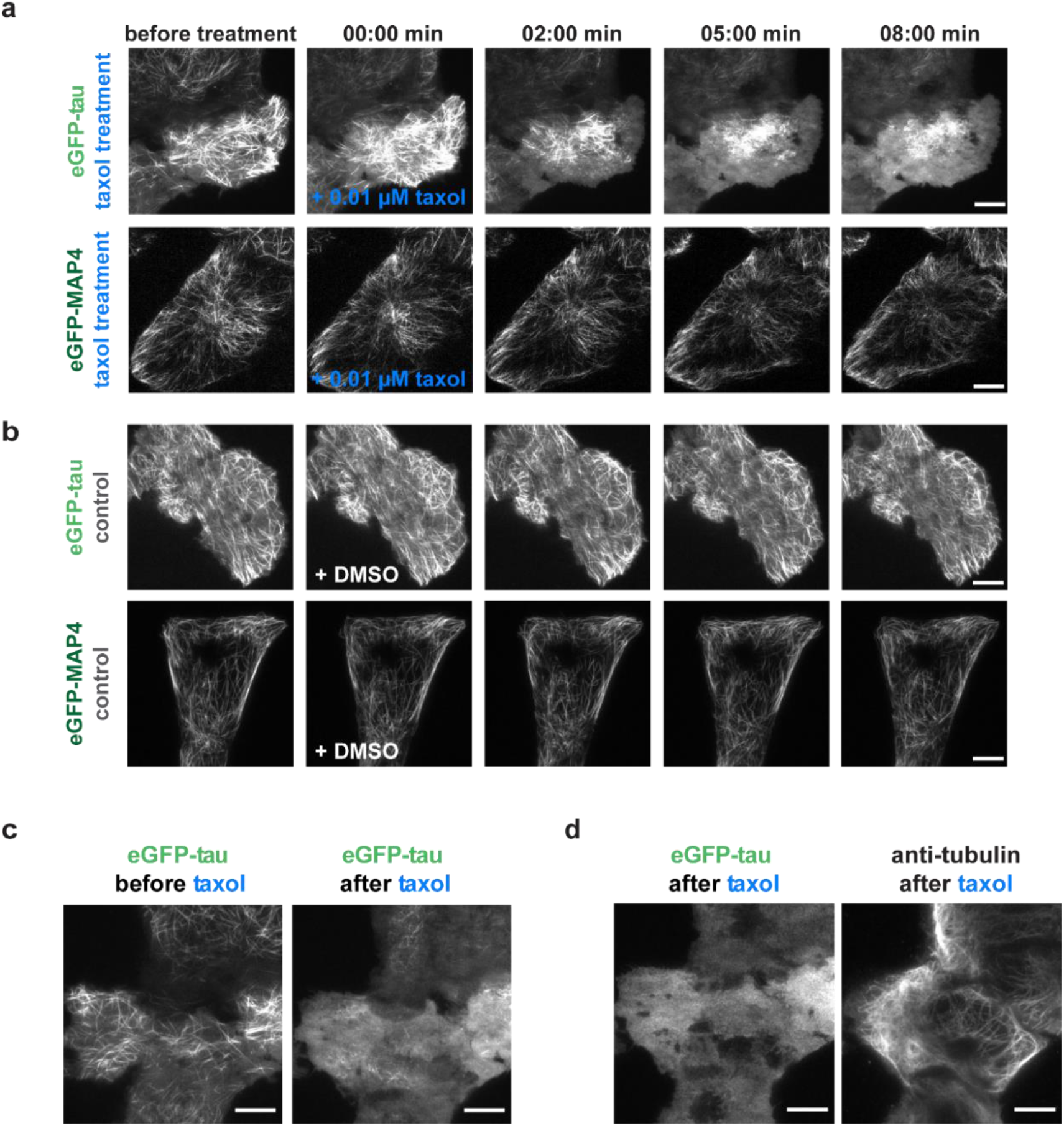
**a.** Time lapse micrographs of U-2 OS cells overexpressing either eGFP-tau (top panels) or eGFP-MAP4 (bottom panels) treated with 0.01 μM taxol. After the addition of taxol, eGFP-tau delocalized from the microtubules, while eGFP-MAP4 remained on the microtubules after the taxol treatment. Scale bars: 10 μm. **b.** Time lapse micrographs of U-2 OS cells overexpressing either eGFP-tau (top panels) or eGFP-MAP4 (bottom panels) treated with DMSO (control) **c.** Fluorescent micrographs from the beginning and end of taxol treatment showing that eGFP-tau delocalized from microtubules. Scale bars: 10μm. **d**. Cells were fixed after taxol treatment, stained against α-tubulin, and imaged by TIRF microscopy showing that microtubules were still present after eGFP-tau delocalization. Scale bars: 10 μm.

## Methods

### Microtubule Assembly

Porcine brains were obtained from a local abattoir and used within ∼4 h of death. Porcine brain tubulin was isolated using the high-molarity PIPES procedure then labeled with biotin NHS ester, Dylight-405 NHS ester, Alexa-647 NHS ester, or Atto-647 ester as described previously (Castoldi & Popov, 2003; Gell et al., 2011; Hyman et al., 1991; Tan et al., 2018). Biotin-labeled tubulin as well as HiLyte647-labeled tubulin were purchased from Cytoskeleton Inc. (T333P and TL670M, respectively).

*Taxol-lattice microtubules (GTP polymerized, then taxol stabilized; stored and imaged in presence of taxol)* were polymerized from 4 mg/ml tubulin for 30 min at 37 °C in BRB80 (80mM PIPES, 1mM EGTA, 1mM MgCl_2_, pH 6.9) supplemented with 4 mM MgCl_2_, 5% DMSO, and 1mM GTP (Jena Bioscience, NU-1012). The polymerized microtubules were diluted in BRB80T (BRB80 supplemented with 10μM taxol (paclitaxel)) and centrifuged for 30 min at 18000 x g in a Microfuge 18 Centrifuge (Beckman Coulter). After centrifugation the pellet was resuspended in BRB80T.

*GMPCPP-lattice microtubules (GMPCPP polymerized)* were polymerized from 4 mg/ml tubulin for 2 h at 37 °C in BRB80 supplemented with 1mM MgCl_2_ and 1mM GMPCPP (Jena Bioscience, NU-405). The polymerized microtubules were centrifuged for 30 min at 18000 x g in a Microfuge 18 Centrifuge (Beckman Coulter). After centrifugation the pellet was resuspended in BRB80T.

*GDP-lattice microtubules (GTP polymerized, GMPCPP-capped)* were polymerized from 4 mg/ml tubulin for 30 min at 37 °C in BRB80 (80mM PIPES, 1mM EGTA, 1mM MgCl_2_, pH 6.9) supplemented with 4 mM MgCl_2_, 5% DMSO, and 1mM GTP (Jena Bioscience, NU-1012). The polymerized microtubules were centrifuged for 30 min at 18000 x g in a Microfuge 18 Centrifuge (Beckman Coulter). After centrifugation, the pellet was resuspended and incubated for 20 min at 37 °C in BRB80 supplemented with 100 mM MgCl_2_, 10mM GMPCPP and 0.25 mg/ml tubulin for cap formation.

*Speckled microtubules* were polymerized as taxol-lattice microtubules prepared from 4 mg/ml tubulin comprised of 2% biotin-labeled tubulin and 0.133% Atto-647-labeled tubulin.

### Protein Constructs and Purification

For *in vitro* experiments comparing tau-family proteins tau, and MAP2c constructs were cloned into pET28A vectors using Gibson assembly. Full-length human MAP4 (isoform 1) was codon optimized for insect cell expression by Epoch Biosciences before cloning into the pFastbac vector. All constructs contain an N-terminal cassette consisting of a 6xHis-tag, tandem Strep-tags, and fluorophore or SNAPf tag connected by a GS-linker as previously described (Tan et al., 2019). *Mus musculus* MAP2c was acquired from the Kassandra Ori-McKenney lab (Monroy et al., 2020).

For *in vitro* experiments comparing tau-family proteins, tau and MAP2c were expressed and purified as described previously (Hernández-Vega et al., 2017; Tan et al., 2019). Briefly, proteins were expressed using BL21(DE3) cells (Agilent). The cells were grown at 37°C until an optical density at 600nm of .6 then induced with 0.2mM isopropyl-Beta-D-thiogalactoside overnight at 18° C. Cells were resuspended in protein buffer pH 8 (50 mM Tris-Cl pH 8, 2 mM MgCl_2_, 1 mM EGTA and 10% glycerol) supplemented with 1 mM DTT, 1 mM PMSF, and 1:1000 Protease Inhibitor cocktail, and 1% triton-X. The cells were then lysed using an Emulsiflex C-3 (Avestin) before high-speed centrifugation to remove cell debris and unlysed cells. Proteins were then purified using Strep XT Beads (IBA). Tau constructs were further purified by cation exchange using a HiTrap HP SP or MonoS column (Cytiva) in protein buffer pH 7.5 with a salt gradient from 100 mM to 400 mM. Fractions were collected, aliquoted, and flash frozen until the day of experiments.

MAP4 was expressed in insect cells using the Bac-to-Bac system (Thermo Fisher). Cells were infected at ∼2 million cells/ml for 60 hours before harvesting. Cells were resuspended in lysis buffer (50mM Tris pH8, 150mM K-acetate, 2mM MgSO_4_ , 1mM EGTA, 10% glycerol) with protease inhibitor, 1mM DTT, 1mM PMSF, 1% Triton X-100, and DNaseI, and dounced on ice. Cell lysate, after douncing, was cleared by centrifugation at 14,000 xg for 20 min, and loaded onto Streptactin Superflow resin (Qiagen) and extensively washed with lysis buffer. Bound proteins were eluted with 3mM desthiobion (sigma) in lysis buffer. Eluted proteins were loaded on cation exchange column 5ml HiTrapS (GE Healthcare) in lysis buffer pH7.5 and eluted with 0-0.6M NaCl gradient over 40 column volumns. Fractions were collected, concentrated, and flash frozen in LN_2_.

SNAPf-tagged MAP proteins were labeled by incubation with 2-5 μM SNAP dye at 4°C for ∼ 2-4 hours. Unbound dye was removed by passage through a HiTrap desalting column equilibrated in GF150 buffer (25mM HEPES pH 7.4, 150mM KCl, 1mM MgCL_2_). Protein was concentrated using Amicon Ultra concentrators (Millipore) and flash frozen in liquid nitrogen and stored at -80°C.

Dynein-dynactin-Hook3 complexes were purified from rat brain lysate as described previously (McKenney et al., 2014). In brief, the SNAPf-Hook3 (1-552) adapter protein construct was purified by Strep-tag affinity as above and further purified by size-exclusion chromatography using a Superose 6 10/300 column in 60 mM HEPES pH 7.4, 50 mM potassium acetate, 2 mM MgCl2, 1 mM EGTA, and 10% glycerol (McKenney et al., 2014). Dynein-dynactin-adaptor complexes were labelled with 2 µM SNAP-TMR dye (NEB) during the isolation procedure from rat brain lysate (McKenney et al., 2014), and were frozen in small aliquots and stored at -80° C. Protein concentrations were assessed using a Nanodrop One (ThermoFisher). Protein concentrations were given as the concentration of fluorophore (monomer) in the assay chamber.

A truncated, constitutively dimerized Kif1A construct (Monroy et al., 2020), and kinesin-1 (Henrichs et al., 2020) were purified as previously described.

### TIRF microscopy

Total internal reflection fluorescence (TIRF) microscopy experiments were performed on an inverted microscope (Nikon-Ti E, Nikon-Ti2 E) equipped with 60x or 100x NA 1.49 oil immersion objectives (Apo TIRF or SR Apo TIRF, respectively, Nikon) and either Hamamatsu Orca Flash 4.0 sCMOS or PRIME BSI (Teledyne Photometrics) cameras. If necessary, an additional 1.5x magnifying tube lens was used. Microtubules and MAPs were visualized sequentially by switching between microscope filter cubes for Cy5, TRITC and FITC channels or by using a quad band set filter (405/488/561/640). The imaging setup was controlled by NIS Elements software (Nikon). *In vitro* experiments comparing multiple MAPs and motors were performed on either of two custom-built through-the-objective TIRF microscope based on a Nikon Ti-E or Ti-2 microscope body, motorized ASI or Nikon stage, quad-band filter cube (Chroma), laser launch (100 mW, 405 nm; 150 mW, 488 nm; 100 mW, 560 nm; 100 mW, 642 nm), EMCCD camera (iXon Ultra 897) and a high-speed filter wheel (Finger Lakes Instruments) (Tan et al., 2019). All imaging was performed using a ×100 1.45 NA objective (Nikon) and the ×1.5 tube lens setting on the Ti-E or Ti-2. The microscopes were controlled using Micro-Manager 1.4 or Nikon Elements software. All experiments were conducted at room temperature, but the sample chamber was kept at 25°C using an objective warmer (Oko Labs).

TIRF chambers were assembled from acid-washed coverslips as described previously (http://labs.bio.unc.edu/Salmon/protocolscoverslippreps.html) and double-sided sticky tape. Chambers were first incubated with 0.5 mg ml^−1^ PLL-PEG-biotin (Surface Solutions) for 10 min, followed by 0.5 mg ml^−1^ streptavidin for 5 min. MTs were diluted into BC Buffer (80 mM PIPES pH 6.8, 1 mM MgCl2, 1 mM EGTA, 1 mg ml^−1^ BSA, 1 mg ml^−1^ casein and 10 μM taxol) then incubated in the chamber and allowed to adhere to the streptavidin-coated surface for 10 min. Unbound MTs were washed away with TIRF assay buffer (60 mM HEPES pH 7.4, 50 mM potassium acetate, 2 mM MgCl_2_, 1 mM EGTA, 10% glycerol, 0.5% Pluronic F-127, 0.1 mg ml^−1^ biotin-BSA, 0.2 mg ml^−1^ κ-casein and 10 μM taxol). Unless otherwise stated, experiments were conducted in imaging buffer (60 mM HEPES pH 7.4, 50 mM potassium acetate, 2 mM MgCl_2_, 1 mM EGTA, 10% glycerol, 0.5% Pluronic F-127, 0.1 mg ml^−1^ biotin-BSA, 0.2 mg ml^−1^ κ-casein, 10 μM taxol, 2 mM Trolox, 2 mM protocatechuic acid, ∼50 nM protocatechuate-3,4-dioxygenase and 2 mM ATP). All experiments were quantified by pooling data from at least two chambers performed on multiple days.

#### FRAP experiments

FRAP experiments were carried out largely as described previously (Tan et al., 2019). For the FRAP experiments, chambers were prepared as described above then sealed at both sides with vacuum grease. Then a pre-bleach image was acquired by averaging 12 consecutive images. Then, 8 regions were bleached (1 empty background, 1 unbleached portion of the MT, 3 envelopes and 3 lattice or 6 regions on the MT in the case of MAP4) at 2% power without scanning; 5 images were taken before stimulation and 91 images were taken after stimulation, all at 1 s intervals. The FRAPed regions were then background subtracted using the empty background region, then normalized to the unbleached portion of the MT, then to the average of the first 12 consecutive images of each region. N is defined as the number of bleached regions.

#### Kinesin-1 on GMPCPP/taxol-lattice MTs

GMPCPP- and taxol-lattice microtubules were immobilized on the coverslip surface. 600 nM tau-mCherry was added to the microtubules prior to addition of kinesin-1-GFP and incubated for at least 2 minutes to ensure envelope formation. After incubation, 60 nM kinesin-1-GFP was added to the microtubules in presence of 600 nM tau, diluted in imaging buffer (50 mM HEPES pH 7.4, 1 mM EGTA, 2 mM MgCl_2_, 75 mM KCl, 10 mM dithiothreitol, 0.02 mg/ml casein, 10 µM taxol, 1 mM Mg–ATP, 20 mM D-glucose, 0.22 mg/ml glucose oxidase and 20 µg/ml catalase). Imaging of kinesin-1-GFP was performed with 20 ms or 50 ms framerate.

#### Lattice expansion TIRF assay

Silanized coverslips were incubated with low anti-β-tubulin antibody concentration (1 μg/ml in PBS) to ensure low binding of microtubules to the coverslip surface. GDP-lattice microtubules were flushed into the measurement chamber and unbound microtubules were removed with BRB80. 20 nM tau-mCherry was added to the microtubules and incubated for 1-2 minutes. After incubation, tau was removed from the measurement chamber by introducing 20 μl of tau imaging buffer either in presence or absence of taxol (50 mM HEPES pH 7.4, 1 mM EGTA, 2 mM MgCl_2_, 75 mM KCl, 10 mM dithiothreitol, 0.02 mg/ml casein, 0 or 10 µM taxol, 1 mM Mg–ATP, 20 mM D-glucose, 0.22 mg/ml glucose oxidase and 20 µg/ml catalase).

#### SiR-tubulin assay

For the data in Fig. 1i, taxol-lattice microtubules were prepared as described above and kept in BRB80 supplemented with 10 μM taxol and 2 μM SiR-tubulin (#SC002, tebu-bio). SiR-tubulin-lattice microtubules were immobilized on the coverslip surface and unbound microtubules were removed with BRB80 supplemented with 10 μM taxol and 2 μM SiR-tubulin. Prior to the experiment, the solution was exchanged by SiR-tubulin imaging buffer (50 mM HEPES pH 7.4, 1 mM EGTA, 2 mM MgCl_2_, 75 mM KCl, 10 mM dithiothreitol, 0.02 mg/ml casein, 2 µM SiR-tubulin, 1 mM Mg– ATP, 20 mM D-glucose, 0.22 mg/ml glucose oxidase and 20 µg/ml catalase). Finally, tau in SiR-tubulin motility buffer was added to the measurement chamber at the final assay concentration stated in the main text.

For the data in Fig. 3d, taxol-lattice 405- and biotin-labeled microtubules were prepared as described above and kept in BRB80 supplement with 10 μM taxol. For each chamber, microtubules were diluted at least 100-fold into TIRF assay buffer (60 mM HEPES pH 7.4, 50 mM potassium acetate, 2 mM MgCl_2_, 1 mM EGTA, 10% glycerol, 0.5% Pluronic F-127, 0.1 mg ml^−1^ biotin-BSA, 0.2 mg ml^−1^ κ-casein and) lacking taxol but including 1 μM SiR-tubulin (Cytoskeleton inc.) and the mixture was allowed to bind to coverslips for 10 min. After this, assay buffer containing the indicated amount of MAP protein and 1 μM SiR-tubulin was flown into the chamber and MAPs were allowed to bind for 10 min before images were acquired.

#### 1,6-hexanediol and envelope enrichment assays

For 1,6-hexanediol experiments, 0.5 nM GFP-MAP2c or GFP-MAP4 was flowed into the chamber and imaged after a 5-minute incubation. Then a solution of 0.5 nM GFP-MAP2c or GFP-MAP4 in 10%-hexanediol in imaging buffer was introduced into the chamber and imaged after a 5-minute incubation. For envelope enrichment assays, 0.5 nM GFP-tau and 0.5 nM mScarlet-tau was incubated in the chamber for 5 minutes, then a mixture of 0.5 nM mScarlet-tau and 0.5nM sfGFP-tau, or sfGFP-MAP2c, or sfGFP-MAP4 was flowed into the chamber and allowed to incubate for 5 minutes. The mScarlet–tau envelopes were used as fiducials for envelope boundaries. Background-subtracted mean intensities were obtained for a line scan along the MT. Each straight and uninterrupted (no MT overlaps) stretch of MT was counted as a single data point. Data points from two different protein preparations of mScarlet–tau envelopes were pooled. Fold enrichment was calculated by dividing each data point for envelope intensity by the average value of associated lattice intensity.

### Optical tweezers experiments

#### Optical trapping assay

Correlative force measurements and microscopy were performed on an optical tweezers setup equipped with confocal fluorescence imaging and microfluidic system (c-Trap, LUMICKS B.V.). The microfluidic chip contains 5 channels through which separate fluids are flushed using a 5-channel air-pressure driven microfluidic system. The microfluidic system was passivated by BSA (0.1% in PBS, #A0281, Sigma) and F127 (1% in PBS) no later than 100 hours before the experiment. The trap stiffness was calibrated using force calibration (0.5.1) in Bluelake (v1.6.11, LUMICKS B.V.) under zero flow condition. The experiments were performed at room temperature.

#### Microtubule lattice compaction (OT)

To study the compaction of the microtubule lattice by tau in the optical tweezers, two streptavidin silica beads (1.12 µm, #SVSIP-10-5, Sperotech) were captured in two traps. The beads were then moved to the microtubule channel (fluorescently labeled taxol-lattice microtubules in imaging buffer: 50 mM HEPES pH 7.4, 1 mM EGTA, 2 mM MgCl_2_, 75 mM KCl, 10 mM dithiothreitol, 0.02 mg/ml casein, 30 µM taxol, 1 mM Mg–ATP, 20 mM D-glucose, 0.22 mg/ml glucose oxidase and 20 µg/ml catalase) where ends of the microtubules were specifically attached to both beads. The construct was then moved to a channel containing imaging buffer where the beads were slowly moved apart until an increase in the force was detected. The microtubule was then kept in a straight but unstretched position and the flow was minimized (0.01-0.02 bar). The construct was then moved into the channel containing 60 nM tau-mCherry in imaging buffer where the compaction of the microtubule lattice was detected by the increase in force concomitant with a decrease in the distance between the two beads. In a control experiment, the construct was moved into a channel containing imaging buffer in the absence of tau (0 nM tau).

#### Envelope disassembly rate

To study the disassembly rate of tau envelopes on microtubules stretched by an external load compared to unstretched microtubules, we attached a microtubule between two beads as described above and moved the construct to a channel containing 60 nM tau-mCherry in imaging buffer. We then kept the microtubule in the tau channel for at least 1 minute for the tau envelopes to form and moved the beads slowly apart until an increase in the force was detected. The microtubule was then kept in a straight position, while the flow was kept constant (0.1 bar). The construct was then moved into the channel containing imaging buffer in absence of tau where the tau envelopes disassembled. In the experiment where an external force was applied to the microtubule, the beads were slowly moved apart until a force was measured of around 40 pN. The force was kept constant by a feedback loop moving the beads further apart while the microtubule relaxed, during the tau envelopes disassembly. In the control experiment (no external force), the microtubule was kept in the straight but unstretched position throughout the experiment.

### CryoEM characterization of microtubule lattice compaction

For CryoEM experiments, 20 µM unlabeled porcine brain tubulin (Cytoskeleton #T240) was polymerized at 37°C for 2 hours in BRB80 (80mM PIPES pH 6.8, 1mM MgCl_2_, 1mM EGTA) supplemented with 3mM GTP. An equal volume of BRB80 supplemented with 10 μM taxol was added, and the polymerized microtubules were left at room temperature overnight. Directly prior to grid preparation, the microtubules were pelleted at 20000 rcf for 10 minutes and resuspended to 3 µM in fresh BRB80.

Microtubules and MAPs were pre-incubated before being added to EM grids. A mixture of 3 μL 3 µM microtubules and 8 μL MAP was incubated at room temperature for 20 minutes. For the microtubule- only dataset, the MAP was replaced with BRB80. Final MAP concentrations were 27 µM tau, 28 μM MAP2c, and 34 µM MAP4. During incubation, R1.2/1.3 Au300 Quantifoil EM grids were glow-discharged for 40 seconds. Then, 4 μL of the microtubule-MAP mixture was applied to the grid in a Vitrobot Mark II (TFS) set to 100% humidity and 22°C. After a 30s wait, the grid was blotted for 4.5s and plunged into liquid ethane. The grids were loaded into a Polara cryo-electron microscope (TFS) operating at 300kV. Images were collected semi-automatically in SerialEM with a pixel size of 1.35Å/px, defocus ranging from -2 to -3 um, and a dose rate of 25e^-1^/px/second for a 1.5 second exposure. ∼40 images were taken for each dataset. Movie stacks were aligned in MotionCorr2 (Zheng et al., 2017). Straight and uninterrupted microtubule segments longer than 2.5 μm were then manually picked in RELION 3.0 (Zivanov et al., 2018), with the coordinates used to crop out each microtubule. In FIJI, each MT segment was then rotated to line up with the Y-axis of the image and the FFT was calculated. The spatial frequency of the peak in intensity of the 4 nm (J_s_) reflection (Chrétien et al., 1996), corresponding to the longitudinal lattice spacing, was measured manually in FIJI.

### Image analysis

Microscopy data were analyzed using ImageJ (FIJI) (Schindelin et al., 2012). For images displayed in figures 3 and 5, the background was subtracted in FIJI using the ‘subtract background’ function with a rolling ball radius of 50 and brightness and contrast settings were modified linearly. In images in which there was substantial drift, the ‘Descriptor-based series registration (2D/3D+T)’ plugin was used in FIJI with interactive brightness and size detections in the MT channel to register the images.

#### Tau density estimation

Tau density on the microtubules was measured In FIJI by drawing a rectangle around the microtubule and measuring the RawIntDen. The rectangle was then moved to an area directly adjacent to the microtubule where no microtubule is present and the RawIntDen was measured again and subtracted from the RawIntDen of the microtubule.

#### Hill coefficient analysis

The tau density (estimated as described above) on taxol-lattice microtubules was plotted against the tau concentration. The Hill coefficient was obtained by fitting this plot using Matlab curve fitting tool using the Hill equation and weights defined as 1/stdev.

#### Growth velocity analysis

Microtubules with different lattices were polymerized as described above. 20 nM tau was added to the measurement chamber and the length of the tau envelopes was measured after 5 minutes of incubation time on the taxol- and GMPCPP lattice microtubules. For the GDP-lattice microtubules, the growth velocity was measured by taking the full length of the GDP-lattice and dividing it by the timepoint at which the entire lattice is covered by a tau envelope.

#### Kinesin run length and landing rate

Kinesin run lengths were measured manually by determining the position at the beginning and the end of the movement using kymographs compiled in FIJI (FIJI *KymographBuilder* plugin). Landing rates were determined by counting the number of traces within 30 seconds on various lengths of microtubules. Traces were considered only when they visually resembled kinesin-1 landings (based on intensity and size).

#### Microtubule lattice compaction

Compaction of the microtubule lattice was measured using speckled microtubules (see above). MAPs were added at t = 30 seconds at concentrations where microtubules would be partially covered with MAP envelopes. In the case of MAP4, 20nM was added. Fluorescent images were taken with 5 second framerate and individual fluorescent speckles were tracked using FIESTA tracking software (v 1.6.0) to obtain the distance between two neighboring speckles. Compaction was measured by averaging the distance between two neighboring speckles before MAPs were added to the microtubules and compared to the distance between the same two speckles after 5 minutes of incubation with the MAPs. Subsequently, fluorescent images of the MAPs were used to correlate the position of the speckles to the position of the envelopes on the microtubule lattice. Compaction of the microtubule lattice between two neighboring speckles on the microtubule lattice were then assigned the corresponding event: envelope region, non-envelope region, not analyzed (both events occurred). In the case of MAP4, due to the lack of envelope formation, no assigning was required and all distances between speckles were pooled into the same category (‘total’).

#### Relative density of tau

In the local bending experiment of GDP-lattice microtubules, the relative density of tau was measured by estimating the tau density (as stated above) in the position of the bend. The density of tau during flow and after tau removal was normalized to the density of tau at the location of the bend before tau was removed.

#### Microtubule lattice compaction OT

The microtubule lattice compaction by tau was detected by the change in distance between the beads using Bluelake v1.6.11 (Lumicks). The distances between the two beads were measured by averaging the distance before adding tau to the microtubule and averaging the distance when maximum compaction was achieved.

#### Analysis of DDH and KIF1A motility

Kymographs of motor motility were analyzed manually to extract velocity and landing rate data. For velocity, entire visible runs were analyzed, including any pauses. For landing rates, the numbers of processive motors in each kymograph were counted manually. Processive runs were counted if the run was at least 3 pixels (∼ 300 nm) long. For pixel-by-pixel intensity correlation between MAPs and motors, average intensity images were generated in each channel using FIJI and analyzed as described previously (Monroy et al., 2020).

### Live-cell experiments

#### Molecular cloning

The sequence of EGFP-tau was subcloned from the vector pSP6 EGFP-tau to the pcDNA4.0/TO mammalian expression vector using BamHI and AscI restriction enzymes. The sequence of the resulting pcDNA4.0/TO_EGFP-tau plasmid was checked using Sanger sequencing. The sequence coding MAP4 gene was amplified from cDNA generated from the U-2 OS cell line. The amplified fragment was then cloned into the pEGFP-C1 mammalian expression vector using the restriction enzymes HindIII and XbaI. We confirmed that the sequence of the amplified gene corresponds to the RefSeq NM_002375.5 with Sanger sequencing. The resulting plasmid was named pEGFP-C1_MAP4.

#### Cell culture and preparation for the live-cell experiments

The U-2 OS cells (ATCC, HTB-96) were maintained in DMEM + GlutaMAX (Sigma-Aldrich, 61965-026) supplemented with 10% FBS at 37°C in 5% CO2. Two days before imaging, the cells were plated onto 4-chamber Glass Bottom Dish (Cellvis, D35C4-20-1.5-N) to achieve 30% confluency. On the following day, the cells were transfected with the mammalian expression vector pcDNA4.0/TO_EGFP-tau or pEGFP-C1_MAP4 using the HP-Xtreme transfection reagent (Sigma-Aldrich, 636624400). The cells were imaged 24h after transfection, with DMEM medium exchanged for FluoroBright DMEM medium (Sigma-Aldrich, A1896701).

#### Live-cell imaging

The movies were acquired using the Nikon H-TIRF microscope equipped with CMOS Hamamatsu ORCA-flash4.0 LT camera and Okolab module for live-cell imaging. The cells were imaged for 1 minute before the addition of taxol or DMSO, with the frame interval being 700 ms. The imaging was then paused, and taxol or DMSO was added to the medium. The final concentration of the taxol was 0.01 μM. The cells were recorded for additional 10 minutes immediately after the addition of taxol or DMSO.

#### In vivo immunofluorescence

After taxol treatment, the recorded cells were fixed directly in a Glass Bottom Dish using 3% Paraformaldehyde in MSB buffer. The cells were then permeabilized with 0.1% Triton Tx-100 and subsequently stained with the DM1A antibody against α-tubulin (Sigma-Aldrich, T9026). The cells were then kept in the MSB buffer and imaged with the TIRF microscope.

#### In vivo image analysis

All the images and movies were processed in FIJI. The two movies resulting from one recording (movie before the addition of taxol or DMSO and movie after the addition of taxol or DMSO) were merged using the „Concatenate“ command. The shift resulting from manipulating the sample when adding taxol or DMSO was corrected with the plugin Template_Matching and function Align_slices in stacks. The kymograph was generated with the KymographBuilder plugin. The coefficient of variation was determined from the standard deviation of the tau or MAP4 fluorescent signal inside the cell before and 10 minutes after taxol or DMSO treatment.

#### Envelope fission frequency and disassembly rate

The fission frequency and disassembly rate of envelopes on microtubules in eGFP-tau cells were measured manually using FIJI. Fissions were manually counted and divided by the length of the microtubule and time. The length of the microtubule or envelope was obtained by the eGFP-tau signal.

### Statistics and reproducibility

All data were collected from at least two independent trials. All of the repeated independent experiments showed similar results. Unless otherwise stated, all data were analyzed manually using ImageJ (FIJI). Graphs were created using GraphPad Prism v.8.0.1 or Matlab R2020b and statistical analyses were also performed using the same software. Major points on graphs represent data means and the error bars represent variation or associated estimates of uncertainty.

### Author Contributions

Valerie Siahaan: Methodology, Formal Analysis, Investigation, Writing, Funding. Ruensern Tan: Conceptualization, Formal Analysis, Investigation. Tereza Humhalova: Methodology, Formal Analysis, Investigation, Funding. Lenka Libusova: Methodology, Supervision, Writing, Funding. Samuel E. Lacey: Methodology, Formal Analysis, Investigation. Tracy Tan: Investigation, Writing – Review and Editing. Mariah Dacy: Investigation, Writing – Review and Editing. Kassandra Ori-McKenney: Conceptualization, Supervision, Writing, Funding, Formal Analysis. Richard McKenney: Conceptualization, Methodology, Supervision, Writing, Funding, Investigation, Formal Analysis. Marcus Braun: Conceptualization, Methodology, Supervision, Writing, Funding. Zdenek Lansky: Conceptualization, Methodology, Supervision, Writing, Funding.

## Acknowledgements

The authors thank members of the MOM lab for feedback and discussion during the project and the Protein Facility of MPI-CBG and V. Váňová for technical support. We also thank Andrew Carter for critical reading of the manuscript, O. Kučera for establishing the optical tweezers assay and J. Sabó for support in obtaining imaging data. This work was supported by Czech Science Foundation grant 19-27477X to ZL and LL and grant 20-04068S to MB. We acknowledge the financial support from the Charles University Grant Agency (GAUK no. 373821 to VS) and the project “Grant Schemes at CU” (reg. no. CZ.02.2.69/0.0/0.0/19_073/0016935) to TH and VS, grants 1R35GM124889 to RJM and 1R35GM133688 to KMOM. KMOM is also supported by the Pew Charitable Trusts grant A19-0406. We acknowledge the institutional support from the CAS (RVO: 86652036), CMS supported by MEYS CR (LM2015043), and the Imaging Methods Core Facility at BIOCEV, an institution supported by the MEYS CR (Large RI Project LM2018129 Czech-BioImaging) and ERDF (project no. CZ.02.1.01/0.0/ 0.0/16_013/0001775) for their support in obtaining imaging data presented in this paper.

